# “*And if you gaze long into an abyss, the abyss gazes also into thee*”: four morphs of Arctic charr adapting to a depth-gradient in Lake Tinnsjøen

**DOI:** 10.1101/817866

**Authors:** Kjartan Østbye, Marius Hagen Hassve, Ana-Maria Tamayo Peris, Mari Hagenlund, Thomas Vogler, Kim Præbel

**Affiliations:** Faculty of Applied Ecology, Agricultural Sciences and Biotechnology, Inland Norway University of Applied Sciences, Campus Evenstad, Anne Evenstadsvei 78, No-2480 Koppang, Norway. E-mail addresses; Department of Biosciences, Centre for Ecological and Evolutionary Synthesis (CEES), University of Oslo, PO Box 1066, Blindern, No-0316 Oslo, Norway; Norwegian College of Fishery Science, Faculty of Biosciences, Fisheries and Economics, University of Tromsø, No-9037 Tromsø, Norway.

**Keywords:** Adaptive radiation, Ecological speciation, Niche specialization, Population divergence, Morphs, Natural selection, Pleistocene ice-age, Microsatellites, MtDNA, *Salvelinus alpinus*

## Abstract

**Background:** The origin of species is a central topic in biology aiming at understanding mechanisms, level and rate of diversification. Ecological speciation is an important driver in adaptive radiation during post-glacial intra-lacustrine niche diversification in fishes. The Arctic charr *Salvelinus alpinus* L. species complex in the Northern hemisphere freshwater systems display huge morphological and life history divergence in lakes with one or several morphs present, thus offering a unique opportunity to address ongoing speciation mechanisms.

We studied Arctic charr in Lake Tinnsjøen by fishing in four nominal lake habitats (pelagial, littoral, shallow-moderate profundal, and deep-profundal habitats) down to 350 meters depth. Research topics addressed were; (1) to illuminate Holarctic phylogeography and lineages colonizing Lake Tinnsjøen, (2) to estimate reproductive isolation of morphs or fish using unbiased methods, and (3) to document eco-morphological and life history trait divergence. Also, we compared Lake Tinnsjøen with four Norwegian outgroup populations of Arctic charr.

**Results:** Four field-assigned morphs were identified in Lake Tinnsjøen; the planktivore morph in all habitats except deep-profundal, the dwarf morph in shallow-moderate profundal, the piscivore morph in shallow-moderate profundal (less in littoral and deep-profundal), and an undescribed new morph – the abyssal morph in the deep-profundal only. The morphs displayed extensive life history variation based on age and size patterns. A moderate to high concordance was observed between field-assigned morphs and four unbiased genetic clusters obtained from microsatellite variation. MtDNA suggested the occurrence of two minor endemic clades in Lake Tinnsjøen likely originating from one widespread colonizing clade in the Holarctic. All morphs were genetically differentiated at microsatellites (F_ST_: 0.12-0.20; with some ongoing gene flow among morphs, and for most mtDNA comparisons (F_ST_: 0.04-0.38). Analyses of Norwegian outgroup lakes implied colonization from a river system below Lake Tinnsjøen.

**Conclusion:** Our findings suggest post-glacial adaptive radiation of one colonizing mtDNA lineage with divergent niche specialization along a depth-temperature-productivity-pressure gradient. Concordance between reproductive isolation and the realized habitat of the morphs imply that ecological speciation may be the mechanism of divergence. Particularly novel is the extensive morph diversification with depth into the often unexplored deep-water profundal habitat, suggesting we may have systematically underestimated biodiversity present in lakes.

## Background

Revealing processes behind adaptive diversity, and formation of species, are central themes in evolutionary biology. Although studied for a long time, the underlying mechanisms for adaptive radiation and speciation often appear enigmatic. However, our consensus understanding is that adaptive radiation by natural selection has been important in the massive origin of populations and species adapting to various environments [e.g. 1–3].

Scientist continuously search for ideal study systems and species groups, to illuminate how speciation processes are acting under evolutionary scenarios and time-scales. Here, highly recognized model species used as rewarding looking-glasses into the species-formation process comprise e.g. Darwińs finches on the Galapagos Islands, European-Mediterranean sparrows, the *Anolis* lizards, Cichlid fishes, the Threespined stickleback, and Sun-flowers [4–9]. The polymorphic northern freshwater fishes of *Coregonus* and *Salvelinus* species complexes are becoming increasingly recognized as good model-systems in this regard [10–23]. Speciation is a complex issue [e.g. 24], where a supportive theoretical framework presents a plethora of mechanisms and avenues for adaptive diversification towards speciation [e.g. 25–30]. Here, it is important to search for and document adaptive divergence in nature e.g. beyond the littoral-pelagic axis in polymorphic species of *Salvelinus*, to derive at a better understanding of the extent and mechanisms of adaptive differentiation. Across examples of adaptive radiation, similarities exist for patterns and processes, where one could tailor models specifically to each species-system to derive a more thorough understanding of the underlying driving mechanisms by empirically parameterizing theoretical models [31–32]. The insight from combined theoretical-empirical analyses can point to important areas where we need to fill knowledge gaps that surface through predictive theoretical models when attempting to add empirical values.

In the ice-covered northern Eurasian hemisphere, the late Pleistocene ice sheet set the frame for colonization and post-glacial adaptation to lakes as the maximum extent of the ice sheet occurred at ca. 21 000 years before present (ybp) and deglaciation at ca. 10-20 000 ybp [33–37]. The Pleistocene (or Quaternary) ice age started ca. 2.58 million years before present, with several alternating phases of glaciation (of roughly 70 000-100 000 years duration) and interglacials (10 000 - 30 000 years duration) [33, 38–39]. Thus, the Pleistocene ice age dynamics represents a long time series where flora and fauna likely repeatedly colonized new land and retracted to glacial refugia over vast geographical areas. Such conditions created opportunities for allopatric differentiation, secondary contact and sympatric diversification between and within species [40–43]. Thus, Holarctic lakes comprise a unique window into the adaptive diversification process of colonizing Arctic charr (*Salvelinus alpinus*, L) where the degree and rate of novel, or parallel adaptations, can be studied by contrasting old versus young glacial geological systems represented by genetic lineages or carbon-isotope dated lakes. Ecological opportunity for diversification via intraspecific competition and niche radiation in species-poor post-glacial lakes may be an important mechanism in morph- and species formation in several fish taxa [16, 28, 44, 45]. One mechanism that could build up reproductive isolation as a secondary product is termed ecological speciation [46–48], and could have been central in adaptive proliferation trajectories of morphs into all lake niches. With regard to sympatric Arctic char morphs, several evolutionary scenarios may be hypothesized [see also 28]. First, the lake could have been colonized by already divergent genetic lineages (associated with different morphs) coming into secondary contact only after separation for thousands of years in glacial refugia. Secondly, sympatric morphs may represent a real intra-lake sympatric adaptive diversification after colonization of one genetic lineage (comprising one initial ancestral morph). Thirdly, a combination of such scenarios could have occurred, generating temporal dynamics in gene-pool-sharing via expansion-contraction, adaptive divergence, speciation reversal, introgression and hybrid swarm dynamics, and subsequent divergence based on novel combinations of genetic variants to be selected upon. Under such adaptive diversification mechanisms, a set of additional interacting mechanisms may be important such as genetic drift and phenotypic plasticity [28, 49, 50].

The highly polymorphic Arctic charr species complex has a Holarctic distribution and is one of the most cold adapted northern freshwater fish species, with some populations having anadromous life history, while most populations are stationary in freshwater [19, 51, 52]. Arctic charr often occupy species-poor Holarctic lakes, suggesting ecological opportunity for adaptive radiation into available niches [15, 19, 53]. Many Arctic charr lakes apparently only harbor a generalist morph, supported by the relative few studies revealing polymorphism. Some of these monomorphic populations, with a generalist morph, utilize both littoral and pelagial habitats through ontogenetic habitat shifts [19]. In a much fewer set of lakes, two more or less distinct morphs, e.g. a littoral and a pelagic morph, may co-occur [55, 56], suggesting lake-specific temporal persistence of niches for the evolution and coexistence of two different morphs. In a very few lakes, a third morph are found in the profundal, termed the profundal morph, coexisting with e.g. the littoral and pelagic morph [57]. Only in one single lake worldwide, namely Lake Thingvallavatn in Iceland, a set of four sympatric morphs are reported that have radiated into all lake niches; a small and large benthic morph, a pelagic morph and a piscivore morph [e.g. 58, 59]. Arctic charr morphs that adapt to divergent niches may show parallelism among lakes with independent origin of morph-pairs [19, 60]. Here, similar morphs can evolve through parallel- or non-parallel evolutionary routes revealing similar gene expression as seen in independently derived morph replicates of two genetic lineages (Atlantic and Siberian lineage) in Arctic charr [23]. This suggest the presence of a highly robust adaptive system in the Arctic charr complex for deriving the same evolutionary outcome from different genetic starting points (historical contingency: adaptive standing genetic diversity, genomic architecture) as response to similar selection pressures. However, there are often lake-specific differences in morph variance regarding e.g. niche occupation, phenotype, and life history [15, 19, 61, 62]. This large-scale parallel evolution in Holarctic lakes, with similar morphs appearing apparently due to similar selection pressures exerted in same niches, is a unique feature when studying natural selection and early steps in the speciation continuum, making the Arctic charr species complex an excellent model system in evolutionary biology and eco-evo-devo studies.

In our study, we report on a new system harboring a striking diversity in phenotypes and life history, apparently associated with a depth-temperature-productivity-pressure gradient in the deep oligotrophic Lake Tinnsjøen in Norway. The history before our study is as follows. On 20th of February 1944, in the occupied Norway during the Second World War, the Norwegian partisans sunk the railway ferry *D/F Hydro* carrying an estimated 20 barrels with 500 kilo of heavy water (D_2_O) in Lake Tinnsjøen. The German occupation government had the purpose to construct an atomic bomb back home in Germany using D_2_O [63, 64]. It has been debated whether this second world war famous sabotage action hampered or stopped Hitleŕs attempt to produce the atomic bomb. Almost 50-60 years later, in 1993 and 2004, a Norwegian team on their search for the sunken ferry, making a second world war news report regarding the presence of heavy water on the ferry, was able to locate it at 430 meter depth using a ROV-submarine, but also at the same time observed small fish residing at the bottom. That team successfully retrieved two fish specimens that was later classified as Arctic charr [65]. The knowledge about the Arctic charr diversity within Lake Tinnsjøen up to that date comprised a study by Hindar et al. [66] showing that a dwarf and planktivore morph grouped together (being statistically different from each other) compared to yet other Norwegian lakes when analyzing allozymes. From old age, local fishermen in Lake Tinnsjøen have recognized a rare deep-water morph of Arctic charr locally named “Gautefisk” (“Gaute” is a Norwegian male name, and “fisk” is fish in Norwegian). This morph has different coloration from other morphs in the lake, and different body proportion, weighting up to 4-6 kg [67]. Thus, when summarizing available information, a set of four morphs were suggested in Lake Tinnsjøen.

As no progress occurred considering scientific studies on the small white fish from the bottom of the lake from the ROV team and associated researchers, we decided to perform a fish survey ourselves that was conducted in the lake in 2013 to document the occurrence of morphs. We set up three main research topics with regard to the Lake Tinnsjøen Arctic charr diversity; (1) to illuminate the phylogeography and ancestral lineages colonizing Lake Tinnsjøen (mtDNA-CytB sequences), (2) to estimate reproductive isolation of field assigned morphs or fish assessed using unbiased methods (microsatellites), and (3) to document eco-morphological and life history trait divergence (body-shape, proportional catch in habitat, age). To accomplish these tasks we collected fish in different habitats in the pelagial, littoral, shallow-moderate profundal and in the deep-profundal. In the field, we classified fish to morphs from exterior phenotype, while in laboratory we assessed morphological (body shape) and genetic divergence using mtDNA and nDNA markers. We further performed a Holarctic phylogeography with online genetic sequences to evaluate lineages colonizing Lake Tinnsjøen. The strength of association of field-assigned morphs and genetically identified morphs using microsatellites (i.e. genetic clusters) was tested. We compared mtDNA and nDNA in Lake Tinnsjøen with a set of four Norwegian outgroup lakes. Using a putative ancestor below in the same drainage, we compared body shape to the Lake Tinnsjøen morphs.

## Methods

### Material used for different analyses

The material used for various analyses is summarized in Additional file: Table S1.

### Study area, fish sampling and field-assigned morphs

Lake Tinnsjøen (60 38 15.6 North, 11 07 15.2 East) is a long (35 km), large (51.38 km^2^) and deep (max depth of 460 m, 190 m mean depth) oligotrophic lake in southeastern Norway (Fig. 1a, b) [68]. High mountain sides surround the lake descending steeply into the lake resulting in a relative small littoral area compared to an extensive pelagic volume and a large profundal area. In the southern and northern end of the lake, larger littoral areas exist with shallow depths. The littoral zone is exposed to the elements such as wind and waves. The shoreline is monotonous with few bays and only one small island. The littoral zone is composed mostly of bedrock, large boulders, smaller rocks as well as sand in less exposed areas and in the deeper layers. The pelagic zone is extensive. The profundal appears to differ structurally in shallow and deep areas - composed of bedrock, boulders, sand and larger-sized organic matter in shallow areas while more fine particulate organic detritus dominates in the deep-profundal areas (based on organic matter on catch equipment and from videos by the Norwegian Broadcasting Company (www-link, no longer valid)). A survey in Lake Tinnsjøen in June 2006 by Boehrer et al. [69] gave an oxygen concentration of 11.5-12.0 mg/l from surface down to 460 m depth, a temperature profile from 4.0-3.3 °C from 50-460 meters depth, conductivity of 10.0-8.0 μS/cm from 0-460 m depth, and dissolved oxygen ranged 90-85% from 0-460 m depth. Lund [70] sampled Lake Tinnsjøen once a month from December 1946 to December 1947 and found that below ca 80 m depth the temperature was at a constant 4 °C (depth stratified), while warming up to ca. 18-20°C in top layer in summer. Thus, Lake Tinnsjøen likely offers a divergent temperature profile (as well as light, pressure and productivity) among habitats, depths and niches, along pelagic and littoral-benthic depth-gradients from surface to 460 m.

**Fig. 1.**
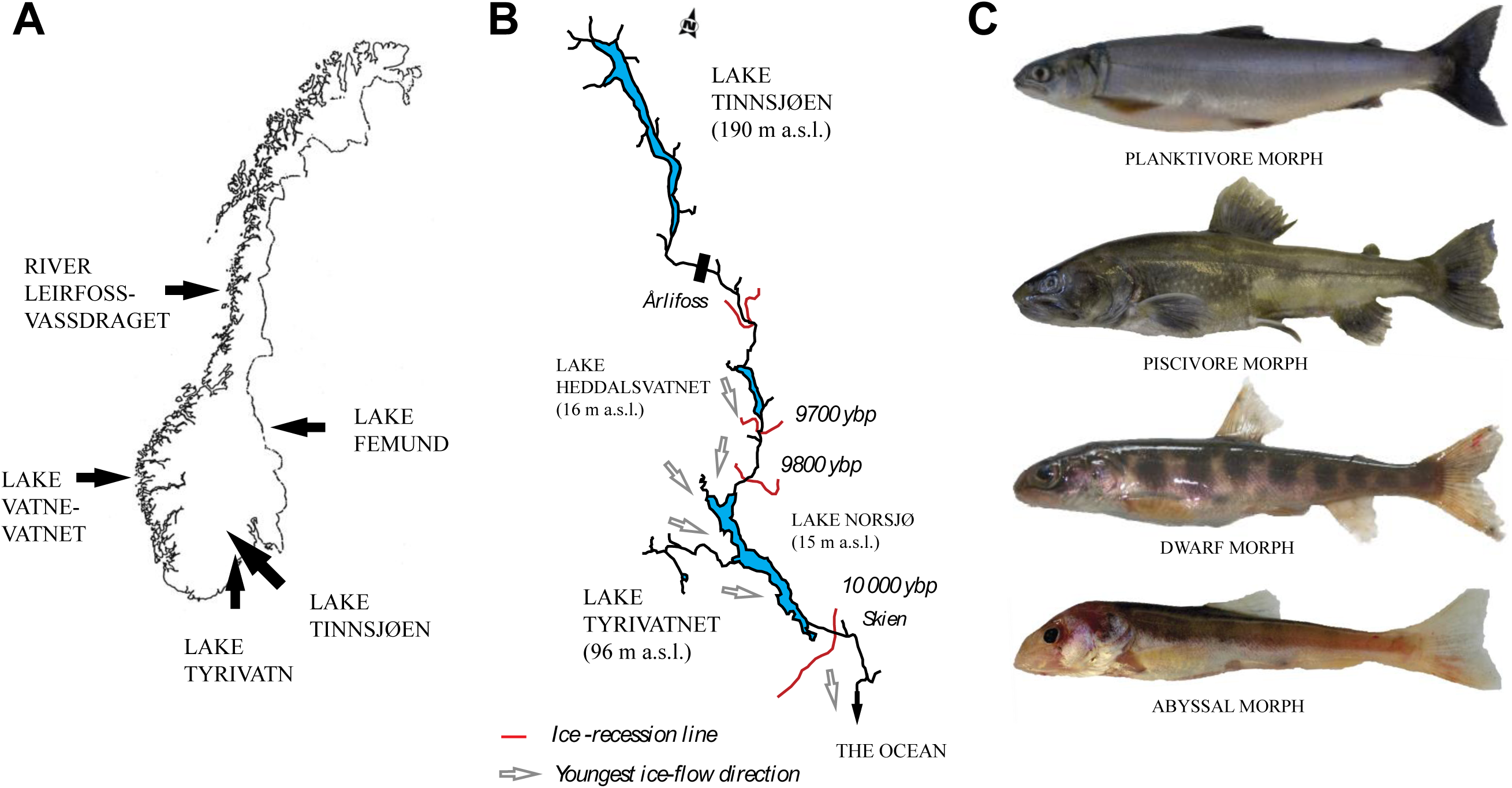
**(A)** Norway with Lake Tinnsjøen and the four outgroups sampled. (**B**) The River Skiensvassdraget wherein Lake Tinnsjøen is situated. Red lines denote dated ice-recession lines in years before present (ybp) based on Bergstrøm [105]. Grey arrows denote the youngest ice-flow direction in the end of the Pleistocene glaciation based on Bergstrøm [105]. The black bar at indicates the upper deposits of marine sediments. (**C**) The four nominal field assigned arctic char morphs (FA-morphs) observed within Lake Tinnsjøen (note: fish scaled to the same length).

We collected Arctic charr from Lake Tinnsjøen during 2013 and from four additional outgroup populations (see below) North, West, East and South of Lake Tinnsjøen in 2013-2015 (Fig. 1a). Fish were caught in four lake habitats (can be viewed as crude nominal niches for individuals and morphs) in Lake Tinnsjøen using equipment described below. At this stage, we do not reveal the exact sampling sites until the taxonomic status of the new abyssal morph has been described and conservation biology authorities in Norway have considered the situation with regard to its conservation value. Particularly relevant here is the population size and uniqueness of the new discovered morph, and what conservation status it merits. As the lake have steep mountain sides entering the lake, it is hard to place equipment precisely at predetermined positions. Thus, habitat and depth ranges fished were grouped to be able to compare catch among four nominal lake habitats. Some fish equipment overlapped to some degree with catches across habitas. However, depth measurements were taken at catch location and noted in the field log, which allowed for later reallocation of the catch effort. As such, the lake habitats defined are quite crude categorizations of habitat and depth ranges, but generally fits with limnologic description of lake niches. The four lake habitats (nominal niches) sampled (and defined by us) in Lake Tinnsjøen in 2013 were; (i) the pelagial (gillnets at <20 m depth, in areas with depths of >30 m, and >50 meters from the shore), (ii) the littoral (gillnets from shore <20 m depth), (iii) the shallow-moderate profundal (gillnets, traps and hook and line from shore at >20 m and <150 m depth), and (iv) the deep-profundal (traps at >150 m depth, > 100 m from the shore).

Sampling was conducted with gill-nets, baited anchored longlines, and traps. Initially, we aimed at fishing with a standardized effort **x** equipment in all niches, but due to the experimental nature of fishing at depths >150 m, and the low fish density, it was difficult to obtain a sufficient sample size. Thus, we intensified the effort in the different habitats with the catch methods that worked best. As such, the material obtained may not be fully representative of fish populations at all depths and habitats, but represents an opportunistic sampling strategy under quite challenging fishing conditions. We used different monofilament series coupled in gangs when fishing with gillnets. In the pelagial, we used a 12-panel multimesh Nordic series (each net; 6 x 60 m) with mesh size (in the following order) of 43, 19.5, 10.0, 55.0, 12.5, 24, 15.5, 35.0, 29.0, 6.3, 5.0 and 10.0 mm (knot to knot), and extended Jensen floating series (each net: 6 x 25 m) with mesh size; 13.5, 16.5, 19.5, 22.5, 26.0, 29.0, 35.0, 39.0, 45.0 and 52.0 mm. In the littoral, we used extended Nordic- and Jensen littoral net series (each net; 1.5 x 60 m or with the same mesh size as in the pelagic zone) including extra nets of some of the largest meshes. We used traps at 20-60 m depth, and Jensen littoral net series (see above for specifications) and hook and line down to 150 m depth in the shallow-moderate profundal. In the deep-profundal we used traps at 100-350 m depth. The baited anchored longlines (ca 220 m long; 3-4 mm line; 180 hooks; size 1, 1/0 and 2), aimed at catching piscivorous Arctic charr, were placed vertically mostly close to the shoreline (<100m) and in a few cases horizontally at the bottom. These attempts resulted in a low catch, and thus the hook and line approach was not used extensively. Nets and baited lines where checked after 12 hours, traps could be out for 48 hours. A motorized winch was used for hauling equipment. All catch was grouped in lake habitats (nominal niches) despite different types of gear used. A total effort of 42 Nordic multimesh and 225 Jensen - net nights, 1001 trap nights, and 27 line nights were implemented in fishing. Besides Arctic charr, we caught brown trout, perch (*Perca fluviatilis*) and minnows (*Phoxinus phoxinus*) (catch statistics not reported as being minute, ie. < 10 individuals in few locations). The lake only hold these four fish species. Minnow was introduced into Lake Tinnsjøen recently (likely in the timeframe 1960-1970’s).

Fish were killed using an overdose of benzocain and transported dead on ice to the field laboratory at Lake Tinnsjøen. In the field, all the fish were subjectively assigned into four nominal morphs based on exterior morphology, being; (i) planktivore, (ii) dwarf, (iii) piscivore, and (iv) abyssal (see representative individuals in Fig. 1c).). Each fish was classified as one of the four morphs despite variation within morphs and subsequent uncertainties This field assignment of morphs was labelled as field-assigned morphs (FA-morphs). Length and weight were recorded, with sex and maturity stage, and age from otoliths in the laboratory. A DNA sample was taken in the field and stored on 96% EtOH for use in analyses (see description below).

The four additional outgroup populations of Arctic charr were situated to the North (River Leirfossvassdraget; anadromous sea-running), West (Lake Vatnevatnet), East (Lake Femund) and South (Lake Tyrivatnet), of Lake Tinnsjøen (Fig. 1a). The three latter Arctic charr populations were stationary in freshwater. The sampling equipment, effort and placement varied among lakes comprising gill nets with at least 16.5. 19.5, 22.5. 29.0 mm (knot to knot) and/or modified Jensen series or Nordic multi-mesh panels set in littoral, pelagic, and profundal areas. In the laboratory, these four populations were analysed as described above for Lake Tinnsjøen. A DNA sample was also stored on 96% EtOH for analyses. These four populations were used as selected outgroups in microsatellite analyses, in mtDNA based phylogenetic analyses, and partly in the morphological analyses. Arctic charr in Lake Tyrivatn was inferred as a putative “ancestral state” founder that could have colonized Lake Tinnsjøen, and was thus used for comparative purposes in microsatellite-, mtDNA- and morphometric analyses (Fig. 1a,b). This was anticipated as the lake is situated far below Lake Tinnsjøen in the same watersystem (see supporting argumentation of the most likely colonization routes in the discussion below). Ideally, we would use the real founding population into Lake Tinnsjøen, but this is not known.

### Phylogeography and the ancestral lineages colonizing Lake Tinnsjøen based on mtDNA

DNA was isolated from pectoral fins using the E-Z96 Tissue DNA Kit (Omega Bio-tek) following the manufactures instructions. Quality and quantity of isolated DNA was assessed using a NanoDrop spectrophotometer and agarose gel electrophoresis. A 851 base pair fragment of the mitochondrial DNA (mtDNA) gene for Cytochrome B (CytB) was amplified using a standard primer pair, FishCytB_F (5’ ACCACCGTTGTTATTCAACTACAAGAAC 3’) and TrucCytB_R (5’ CCGACTTCCGGATTACAAGACCG 3’) [71] in 10 µl polymerase chain reactions (PCR). The reactions consisted of 1 µl 10 x PCR-Buffer, 0.3 µl 10 µM dNTP, 0.5 µl of each of the 10 µM F and R primers, 5.5 µl ddH_2_O, 0.2 µl FinnZyme DNAzyme ext Polymerase, and 2 µl DNA template (0.4-0.8 µg). The cycling profile consisted of an initial 5 min denaturation step at 94°C, 32 cycles of 94°C for 30 s, 57°C for 35s, and 70°C for 1 min, followed by a final 10 min elongation step at 70°C. The products were treated with ExoZap^TM^ to remove leftover primers and dNTPs, before running the standard BigDye reaction, using the above primer set in 3.5 µM concentrations. The products were cleaned by precipitation, before sequencing them on an ABI 3130XL Automated Genetic Analyzer (Applied Biosystems), using 80 cm capillaries. All sequences were manually trimmed and verified in Geneious 10 (Biomatters).

For phylogeographical analyses using Cytochrome B, the 851 base pair long sequences were aligned in Mega 7.0.26 using default settings [72]. Sequences were interpreted mostly based on both forward and reverse readings (but in a few cases, only one sequence direction was readable). A set of 115 Norwegian sequences were retrieved where sample size range 21-22 for the four Lake Tinnsjøen FA-morphs and a sample size of 5-9 for the four Norwegian outgroup lakes (Additional file 1: Table S1).

For larger scale comparison of phylogeny, highly similar sequences were retrieved using BLAST (https://blast.ncbi.nlm.nih.gov/Blast.cgi) (Fig. 2a,b, Additional file: Table S2a). A cutoff of 200 highly similar sequences were downloaded from Blast (including various *Salvelinus* taxa), aligned as described above and analyzed with sequences from Lake Tinnsjøen and the four Norwegian outgroup lakes.

**Fig. 2.**
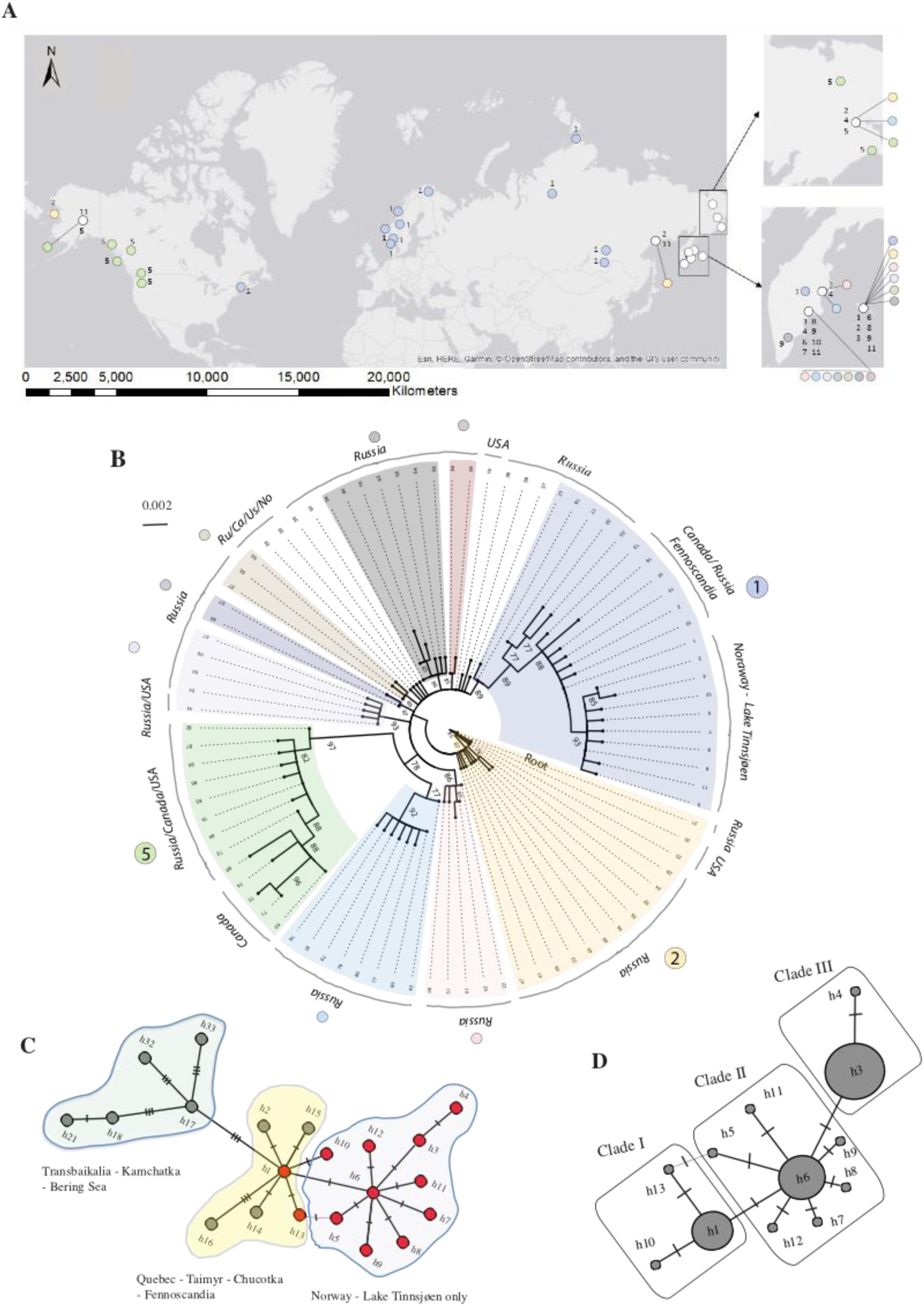
**(A)** Distribution of 88 mtDNA-Cytochrome B mtDNA haplotypes compared with major clades in different colors according to figure B. White circles denote haplotypes not well supported in figure B. **(B)** Circular phylogenetic tree of sequences mapped in figure A. Here, a total of 13 Norwegian sequences and 75 haplotypes retrieved from GenBank (using a cut-off of 200 highly similar BLAST sequences) are compared. Hhere, haplotype 31 was found to be the most ancestral when rooted with three distant salmonid taxa (*Salmo trutta*, *Oncorhynchus kisutch* and *Coregonus lavaretus*) (tree not shown). Major supported clades have different colors. Main geographical regions are named on the outer circle. **(C)** A minimum spanning network of haplotypes (not frequencies) in the major light purple clade comprising Lake Tinnsjøen with geographical areas described. Haplotypes in red was found in Lake Tinnsjøen. **(D)** A minimum spanning network with frequencies for haplotypes in Lake Tinnsjøen. The three clades harbor haplotypes from Lake Tinnsjøen while more clades can be found in the major light purple clade in figure B.

The best substitution model for the combined dataset (115 Norwegian and 200 Blast sequences) was interpreted using online server IQ-Tree (http://www.iqtree.org/) with 10 000 ultrafast bootstrap iterations [73–75]. Here, the best substitution model revealed was TN + F + I [76] (Additional file: Table S3).

A circular phylogenetic tree using the TN + F + I model was visualized in Treview 1.6.6 [77] using all the 88 observed haplotypes from the joint dataset from the 115 Norwegian sequences and 200 Blast sequences. Earlier, in another tree, we initially used three outgroup taxa to reveal the most ancient haplotypes in the charr sequences, being; *Salmo trutta* (GeneBank accession; LT617532.1), *Oncorhynchus kisutch* (KJ740755.1) and *Coregonus lavaretus* (AJ617501.1). This latter tree is not shown, but the most ancestral *Salvelinus* sp. sequence revealed from this analysis presented in results.

A map was made [78] for the joint dataset of the 88 sequences and plotted geographically with regard to 10 selected major clade configurations. Clade definition and selection was done to visualize the large geographical scale patterns of sequences (although alternative clade definitions do exist).

A major large-scale phylogenetic branch including the Lake Tinnsjøen haplotypes were used for drawing a minimum spanning network in PopART (http://popart.otago.ac.nz) [79], when not considering frequencies of haplotypes. This major clade which harbored 21 haplotypes had good statistical support (89%) from the remaining haplotypes and were selected for further resolution, covering a large geographical range. The purpose with this clade selection was to have an in-depth look at the putative radiation and geographical distribution of the closest genetic relatives to the Lake Tinnsjøen morphs.

In addition, for Lake Tinnsjøen, a network was built using TCS v1.21 [80] visualizing only the mtDNA sequences found inside the lake to reveal putative formation of subclades after initial founding colonization.

The three 1-mutational step clades (clade 1-1-, 1-2, and 1-3) revealed in TCA in Lake Tinnsjøen and the four Norwegian outgroup lakes were only analyzed for putative signals of population demographic changes in DnaSP v6.11.01 [81] using pairwise sequence distribution and Tajimàs D and Fu and Lìs D estimators [82, 83].

For the four outgroup lakes, the four FA-morphs and the three mtDNA clades in Lake Tinnsjøen, the number and percentage of haplotypes were calculated along with genetic diversity estimators in DNAsp v6.11.01 [81].

### Reproductive isolation of field assigned morphs or fish assessed using unbiased methods

A set of 11 microsatellites were amplified and analyzed after procedures in Moccetti et al. [62] (Additional file: Table S2b,c). 3-6% negative controls per plate and 4% replicate samples were included in the analysis to control cross-contamination and consistency of genotypes. All negative samples were blank in the fragment analysis and all replicate samples had matching genotypes. The genotypes were scored in Genemapper 3.7 (Applied Biosystems) using automatic binning in predefined allelic bins. All genotypes were subsequently verified by visual inspection independently by two persons.

Deviation from Hardy Weinberg equilibrium (HWE) and linkage disequilibrium (LD [84] was estimated using GENEPOP 4.6 [85, 86] implementing an exact test. The presence of LD may lead to erroneous conclusions if loci do not have independent evolutionary histories. Loci exhibiting significant LD should be excluded from analyses. False discovery rate (FDR) corrections [87] was used to test for significant HWE and LD adjusting p-values for multiple tests. The results showed that out of 40 tests of departures from HWE, significant deviations were not found in any loci or populations after FDR correction. Significant LD was discovered between loci SCO204 and SCO218. Thus, locus, SCO204 was removed, and a total of 10 loci were used in the following genetic analyses.

GENEPOP 4.6 [85, 86] was used to calculate number of alleles, expected and observed heterozygosity, as well as genetic divergence between populations (F_ST_) using log-likelihood based exact tests. The software HP-RARE 1.0 [88] was used to calculate standardized private allelic richness (A_p_) and standardized allelic richness (A_r_) accounting for differences in sample size. A_p_ and A_r_ was calculated with rarefaction using the minimum number of genes in the samples i.e. 28 genes.

The software MICRO-CHECKER 2.2.3 [89] was used to check for null alleles, stutter-errors, large allele dropout and size-independent allelic dropout. Of the ten loci, MICRO-CHECKER found one loci to exhibit homozygote excess, potentially due to null alleles, being SalF56SFU. Due to the presence of null alleles, the program FREENA [90, 91] was run to correct for this using the ENA method (Excluding Null Alleles). The FREENA software was run with 5000 replicates, and corrected F_ST_ values were used.

The software LOSITAN [92, 93] was used to test if loci were under selection. Using loci under selection may give erroneous results of genetic structure and F_ST_ values [93]. All loci were run under both the stepwise mutation model (SMM) and the infinite alleles model (IAM) using 100,000 simulations under the “Force mean F_ST_”, and “Neutral mean F_ST_” alternatives. None of the loci were indicated as candidates for directional selection.

Genetic differentiation (F_ST_) was estimated in GENEPOP 4.6 [86] comparing Lake Tinnsjøen and the four outgroup lakes, the four FA-morphs and the four outgroup lakes, and among revealed genetically defined morphs (termed GA-morphs, with a definition of genetic morphs being q>0.7 based on STRUCTURE results; see details below) in Lake Tinnsjøen.

To determine the most likely number of genetic clusters (K) the software STRUCTURE [94] was run using 500 000 burn-in steps and 500 000 Markov Chain Monte Carlo (MCMC) repetitions with 10 iterations, considered as a high enough number to reach convergence. STRUCTURE was run a first time with the individuals from Lake Tinnsjøen and the four Norwegian outgroups; Lake Femund, Lake Tyrivatnet, Lake Vatnevatnet and Lake Leirfossvassdraget. Secondly, a hierarchical approach was performed where the population that deviated the most from the remainder of the populations was removed, and all remaining populations were run a second time. This was repeated until no more clustering was found. The number of genetic clusters was estimated by calculating the logarithmic probability (LnP(K)) and ΔK which is based on changes in K [95]. The most likely number of clusters was determined using STRUCTURE- HARVESTER [96]. According to reccomendations by Hubisz et al. [97], STRUCTURE was also run with the LOCPRIOR function wich incorporates geographic sampling locations using default values.

Based on K-clusters results from the STRUCTURE analysis, we assigned different genetic populations or morphs in Lake Tinnsjøen (GA-morphs). We further contrasted Lake Tinnsjøen with the four outgroup lakes. Assignment analyses was based on K-clusters of individuals with q-values of >0.7 to its own cluster, evaluated as belonging to this population. Individuals with q-values <0.7 were interpreted as being hybrids of unsure population origin.

As an alternative way to test genetic differentiation, we first conducted a principal component analysis in Genetix 4.05.2 [98] based on microsatellite alleles, revealing the variation explained along the first two axes to be; PC1 (33.0 %) and PC2 (17.3 %). Then, we tested for differentiation among the five lakes for PC1 and PC2 using a nonparametric multiple comparison test (Steel-Dwass all pairs) in JMP 11.2 [99]. In addition, we used the same approach for testing differentiation, but now along PC1-3, for four FA-morphs in Lake Tinnsjøen as described above, by only sub-setting Lake Tinnsjøen from the five lake dataset.

### Eco-morphological and life history trait divergence in the Lake Tinnsjøen charr morphs

In Lake Tinnsjøen, association between habitat occurrence with FA-morphs or GA-morphs was tested using χ^2^ statistic in JMP 11.2 [99]. See bathymetric map in Figure 3a. The main purpose here was to reveal association of morphs (FA or GA) and habitat at catch, however, we are aware of the putative bias in having used different fishing gear in different habitats. See also previous section on fish sampling regarding overall issues related to statistical testing.

**Fig. 3.**
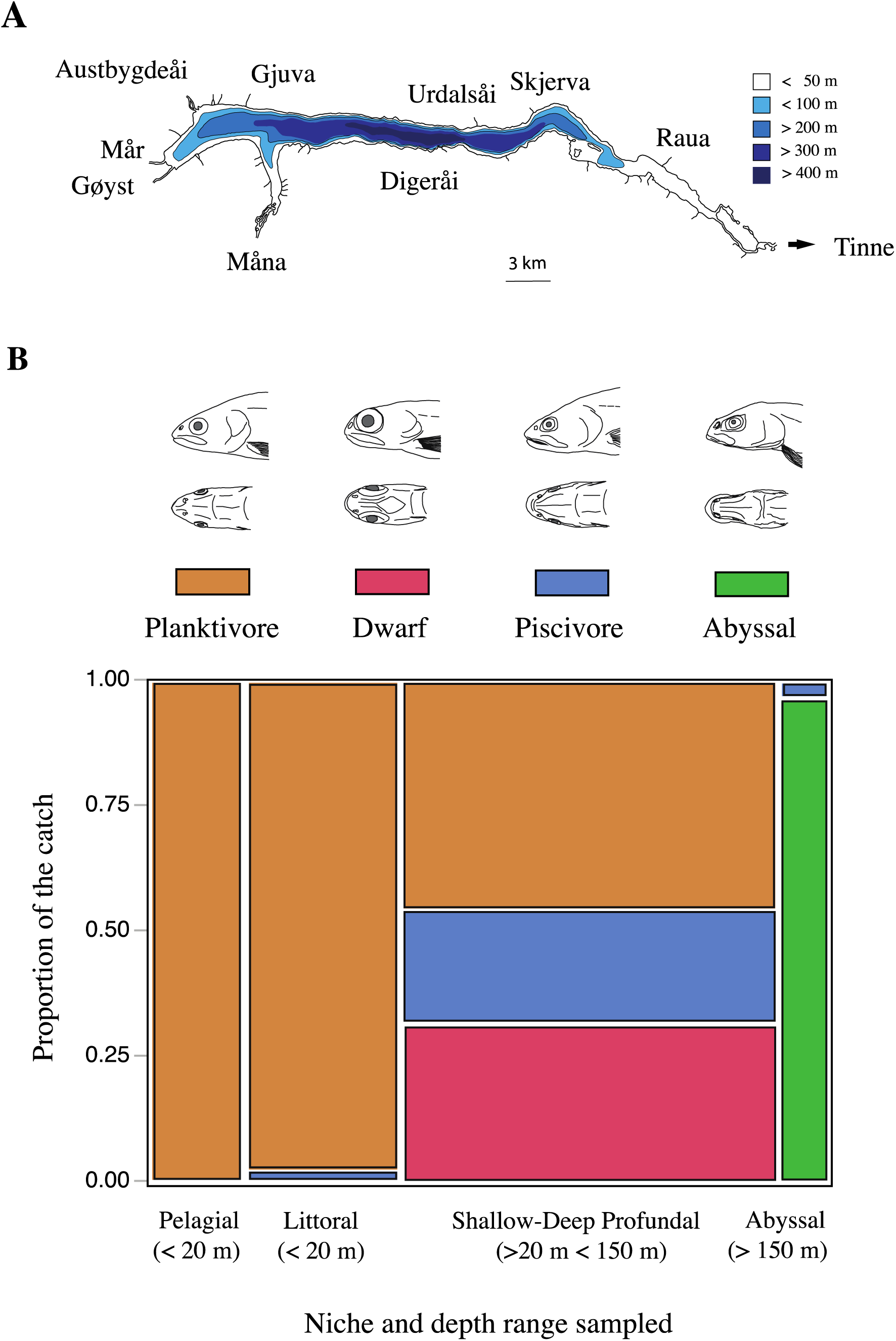
**(A)** A crude bathymetric map of Lake Tinnsjøen (modified from; The Norwegian Water resources and Energy Directorate; http://gis3.nve.no/metadata/tema/DKBok1984/Dybdekart_1984.htm) [68]. (**B**) The association between the catch of the four FA-morphs in the four lake habitats in Lake Tinnsjøen. A drawing of representative heads (lateral and ventral views) of each of the four FA-morphs are given in top panel.

A discriminant analysis in JMP 11.2 [99] was used to test for association between GA-morphs and FA-morphs.

A geometric morphometric analysis using landmarks to reveal body shape was conducted using; Lake Tinnsjøen only, and secondly Lake Tinnsjøen and Lake Tyrivatn in the river drainage to the south of Lake Tinnsjøen. In the latter analysis, the idea was to evaluate the phenotype of the putative ancestral founder that could have colonized Lake Tinnsjøen, and how the Arctic charr in Lake Tyrivatn was morphologically assigned to the FA-morphs in Lake Tinnsjøen.

A Canon EOS 550d mirror reflex camera (Canon lens EFS 18 - 55 mm and macro lens EFS 60 mm; F20 ISO1600 AV, blitz) was used to photograph (JPEG) fish. Photos were taken in a Styrofoam box with a permanent standardized light. Fish were placed in natural position with their left side fronting the camera. All fish which had inflated swim bladders were carefully punctuated so that inflation did not affect body shape (subjective correction).

After digitalization in TpsUtil 1.53 [100], transforming JPEG to tps-files, landmarks were scored in TpsDig2 2.16 [101]. A set of 30 landmarks (real and semi-landmarks) were used to capture the body shape of fish, with main focus on the head region (Fig. 4a). Similar landmarks have been used in other studies, but there are no consensus regarding position or number of landmarks to be used. A transparent film with imposed lines helped setting semi-landmarks. To minimize inter-individual scoring bias, all landmarks where set by one person.

**Fig. 4.**
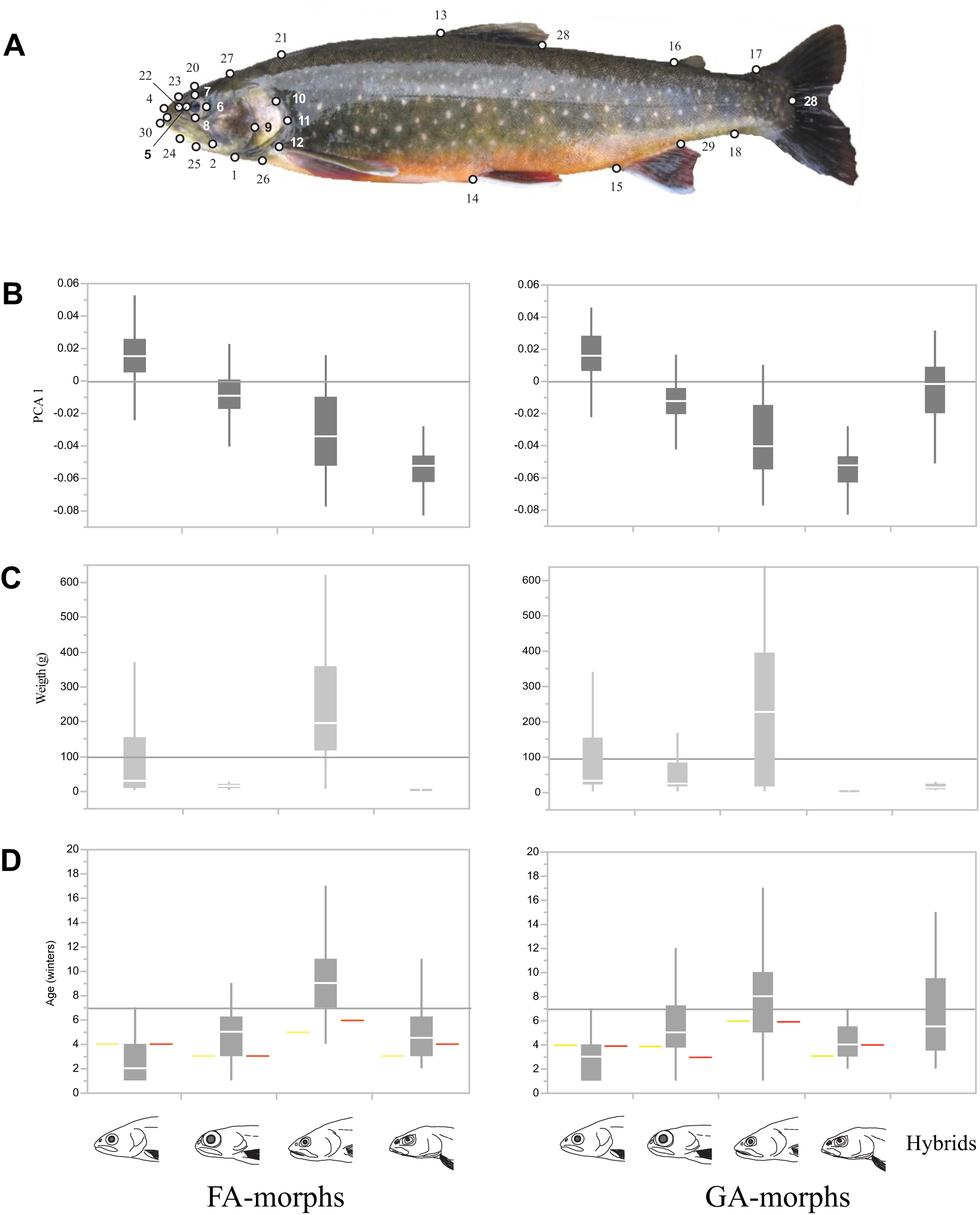
**(A)** The 30 landmarks used for body shape analyses in Lake Tinnsjøen: 1. lower edge of preoperculum, 2. edge of maxillary bone, 3. mouth opening, 4. tip of snout, 5. - 8. eye positions, 9. mid edge of preoperculum, 10. posterior edge of preoperculum, 11. posterior edge of operculum, 12. pectoral fin, 13. and 28.; dorsal fin, 14. pelvic fin, 15. and 29.; anal fin, 16. adipose fin, 17. upper tail root, 18. lower tail root, 19. end of the side line organ, 20. top of head, 21. back above pectoral fin, 22. nostril, 23. over nostril, 24, under-jaw, 25. edge of mouth, 26. lower edge of operculum, 27. transition zone from head to body, and 30. edge of lower lip. **(B)** Principal component axis 1 versus respectively FA-morphs (left panel) and GA-morphs (right panel) based on the 30 landmarks. (**C**) weight versus FA-morphs and GA-morphs. (**D**) Age versus FA-morphs and GA-morphs. The youngest sexually mature male (yellow line) and female (red line) are given. The graphs denote median values (white horizontal line), 25% to 75% (solid blocks), and 10% to 90% percentiles (grey vertical line). In figure **A**-**C** arbitrarily selected horizontal lines have been imposed for helping out visual comparisons among the four FA-morphs and the four GA-morphs, and in two panels compared.

In MorphoJ 1.06 [102], using the TpsDig2 file, extreme outliers were removed from both datasets after an outlier analysis, followed by a Procrustes fit analysis. A principal component analysis with eigenvalues was conducted for each dataset. The first five PC-axes were used for further analyses. For Lake Tinnsjøen, PC-axes 1-5 explained 45-4% of the variation in body shape, with a summed variation of 81.5% (PC1 45%, PC2 14%, PC3 13%, PC4 6% and PC5 4%, respectively). For Lake Tinnsjøen and Lake Tyrivatn, PC-axes 1-5 explained 45-4% of the variation in body shape, with a summed variation of 78.3% (PC1 45%, PC2 13%, PC3 12%, PC4 5% and PC5 4%, respectively). As there were still body length effects on shape after PC-analyses in MorphoJ (likely due to allometric growth), we corrected for body length using a regression of log centroid size on body shape (PC-axes 1-5) in MorphoJ in both datasets, then saving the residuals for further analyses.

To evaluate how concordant body shape was to FA-morphs in Lake Tinnsjøen, we used a discriminant analysis in in JMP 11.2 [99] with linear, common covariance using residuals from the five PC-axes in MorphoJ. Similarly, we tested morphological resemblance in body shape of the FA-morphs with their putative ancestral founder from Lake Tyrivatn. Assignment percentages to each of the categories were recorded for both analyses.

A subset of the fish was used for determining age based on otoliths, immersed in 95% EtOH, read using a microscope [103]. An unfortunate challenge was encountered as the Arctic charr heads had been stored in unbuffered formalin which partly prevented age reading in some fish due to unbuffered formalin eating up parts of the otoliths. However, for age determined fish, we were confident in their age. Some of the morphs had few individuals analyzed for age. Also, it was difficult to determine maturity stage in some fish. This situation prevented a thorough life history analysis. Thus, we present age and body weight distributions revealing the youngest sexually mature male and female (also for body weight distributions).

## Results

### Fish catch and field assigned morphs

A total of 754 fish were caught in Lake Tinnsjøen, comprising 457 Arctic charr, 294 brown trout, 3 perch, and a small number of European minnow (not quantified). In Arctic charr, 63 fish (13.8% of the total catch of Arctic charr) were caught in the pelagial, 105 fish (23.0%) in the littoral, 256 fish (56.0%) in the shallow-moderate profundal, while 33 fish (7.2%) were caught in the deep profundal (Table 1). For brown trout, 101 fish were caught in the pelagial, 131 in the littoral, and 62 in the profundal. European minnow and perch were only caught in the littoral.

**Table 1.**
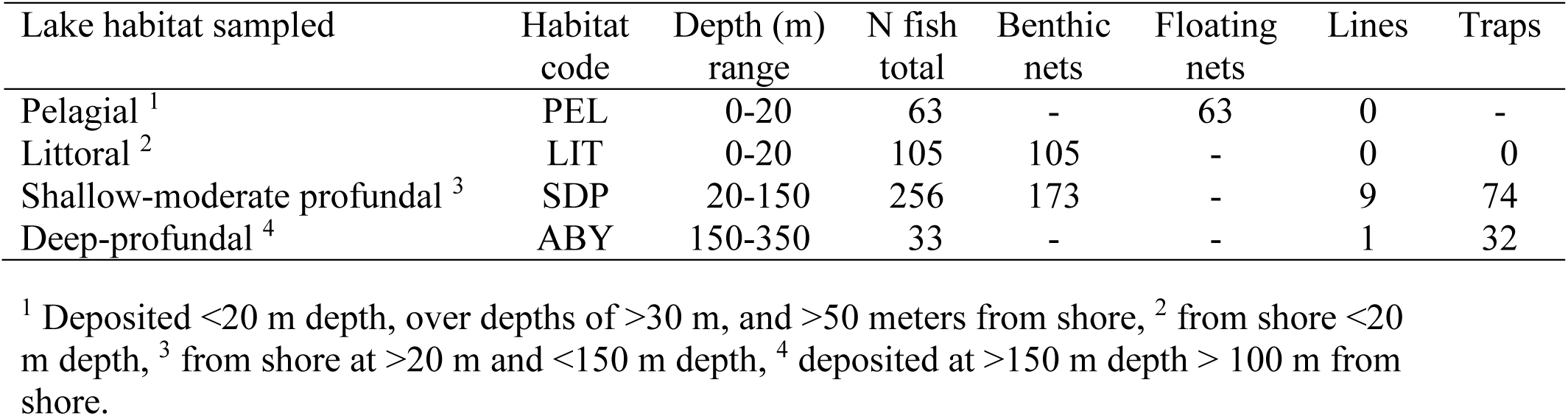
The arctic charr (N=457) collected in Lake Tinnsjøen in 2013 using different equipment. - denote equipment not used in that habitat (niche) while a value of 0 denote equipment used, but no catch in that habitat. The sampling effort was not standardized precluding catch per unit effort.

In Lake Tinnsjøen, the field assigned morphs based on visual appearance (FA-morphs, N=457) revealed 282 fish (61.7%) of the planktivore morph, 81 fish (17.7%) of the dwarf morph, 62 fish (13.6%) of the piscivore morph, and 32 fish (7.0%) of abyssal morph (Table 2).

**Table 2.** Number and catch percentage of the total catch (N=457) partitioned into field assigned morphs (FA-morphs) in the four lake habitats. The bottom row summarize the number and catch percentage in the four habitats across the morphs and the last two columns similarly summarize the catch of the morphs.

| FA-morphs | PEL N | % | LIT N | % | SDP N | % | ABY N | % | In morph | % |
| --- | --- | --- | --- | --- | --- | --- | --- | --- | --- | --- |
| Planktivore | 63 | 13.8 | 102 | 22.3 | 117 | 25.6 | - | - | 282 | 61.7 |
| Dwarf | - | - | 1 | 0.2 | 80 | 17.5 | - | - | 81 | 17.7 |
| Piscivore | - | - | 2 | 0.4 | 59 | 12.9 | 1 | 0.2 | 62 | 13.6 |
| Abyssal | - | - | - | - | - | - | 32 | 7.0 | 32 | 7.0 |
| Across morphs | 63 | 13.8 | 105 | 23.0 | 256 | 56.0 | 33 | 7.2 | 457 |  |

### Phylogeography and the ancestral lineages colonizing Lake Tinnsjøen based on mtDNA

A set of 13 haplotypes (*h1-13*) were found in the combined dataset of Lake Tinnsjøen and the four Norwegian outgroup lakes (Table 3). The 13 haplotype sequences obtained in our study are deposited on GenBank (accession number x-y). Here, 12 of the 13 haplotypes were only found in Lake Tinnsjøen (which lacked *h2*). The four outgroup lakes all had haplotype *h1*, which also occurred in all of the four FA-morphs, while only one outgroup lake, Lake Vatnevatnet, had an additional haplotype *h2*.

**Table 3.**
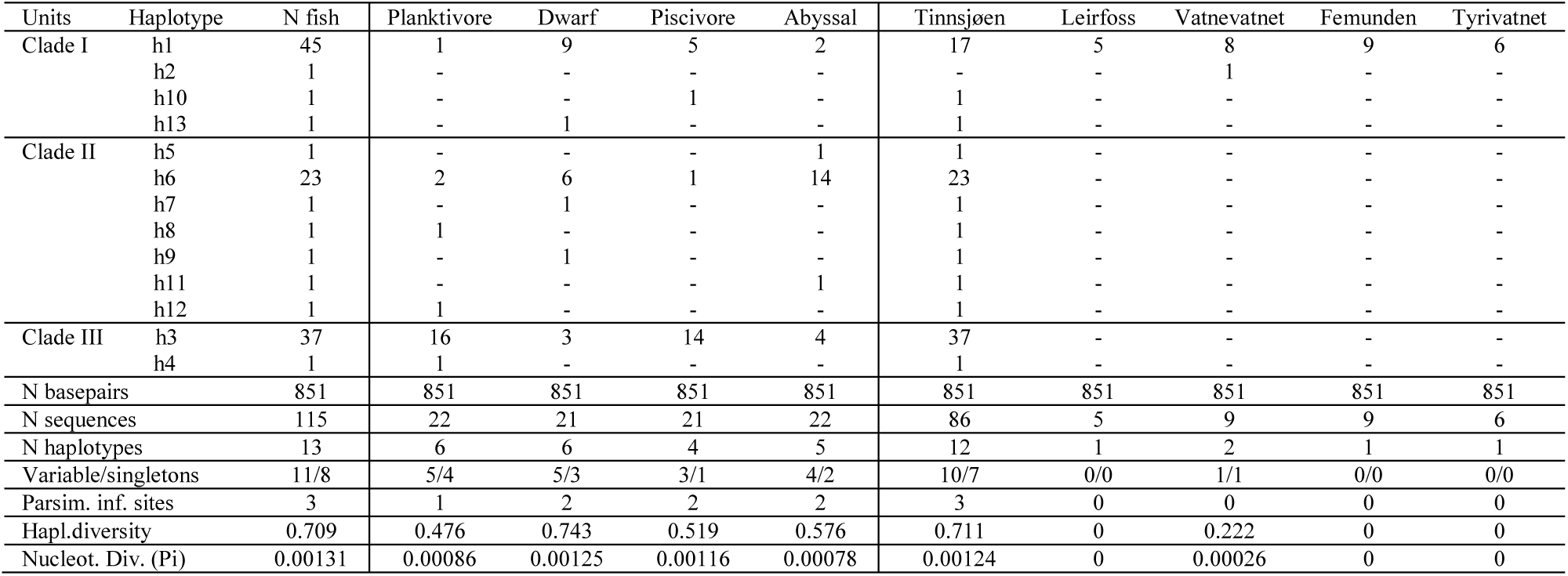
The observed mtDNA-haplotypes in Lake Tinnsjøen and in the four Norwegian outgroup lakes. Colors represents three clades where haplotypes group together in the phylogenetic tree (see Fig. 6). Summarize statistics for genetic variation in the morphs and lakes is also given.

| Units | Haplotype | N fish | Planktivore | Dwarf | Piscivore | Abyssal | Tinnsjøen | Leirfoss | Vatnevatnet | Femunden | Tyrivatnet |
| --- | --- | --- | --- | --- | --- | --- | --- | --- | --- | --- | --- |
| Clade I | h1 | 45 | 1 | 9 | 5 | 2 | 17 | 5 | 8 | 9 | 6 |
|  | h2 | 1 | - | - | - | - | - | - | 1 | - | - |
|  | h10 | 1 | - | - | 1 | - | 1 | - | - | - | - |
|  | h13 | 1 | - | 1 | - | - | 1 | - | - | - | - |
| Clade II | h5 | 1 | - | - | - | 1 | 1 | - | - | - | - |
|  | h6 | 23 | 2 | 6 | 1 | 14 | 23 | - | - | - | - |
|  | h7 | 1 | - | 1 | - | - | 1 | - | - | - | - |
|  | h8 | 1 | 1 | - | - | - | 1 | - | - | - | - |
|  | h9 | 1 | - | 1 | - | - | 1 | - | - | - | - |
|  | h11 | 1 | - | - | - | 1 | 1 | - | - | - | - |
|  | h12 | 1 | 1 | - | - | - | 1 | - | - | - | - |
| Clade III | h3 | 37 | 16 | 3 | 14 | 4 | 37 | - | - | - | - |
|  | h4 | 1 | 1 | - | - | - | 1 | - | - | - | - |
| N basepairs |  | 851 | 851 | 851 | 851 | 851 | 851 | 851 | 851 | 851 | 851 |
| N sequences |  | 115 | 22 | 21 | 21 | 22 | 86 | 5 | 9 | 9 | 6 |
| N haplotypes |  | 13 | 6 | 6 | 4 | 5 | 12 | 1 | 2 | 1 | 1 |
| Variable/singletons |  | 11/8 | 5/4 | 5/3 | 3/1 | 4/2 | 10/7 | 0/0 | 1/1 | 0/0 | 0/0 |
| Parsim. inf. sites |  | 3 | 1 | 2 | 2 | 2 | 3 | 0 | 0 | 0 | 0 |
| Hapl.diversity |  | 0.709 | 0.476 | 0.743 | 0.519 | 0.576 | 0.711 | 0 | 0.222 | 0 | 0 |
| Nucleot. Div. (Pi) |  | 0.00131 | 0.00086 | 0.00125 | 0.00116 | 0.00078 | 0.00124 | 0 | 0.00026 | 0 | 0 |

From the samples in the larger scale phylogeography (Fig. 2a), a total of 75 new haplotypes were retrieved from Blast, comprising 88 haplotypes including the 13 Norwegian haplotypes (Additional file: Table S2). Comparing these 75 haplotypes to the ones found in Norway revealed that only *h1* (in several lakes) and *h13* (in one lake) were found outside Lake Tinnsjøen and the four Norwegian outgroups. Lake Tinnsjøen harbored a set of 10 endemic haplotypes (*h3-h12*).

The major branch in Figure 2b (light purple) including the Lake Tinnsjøen haplotypes were used for drawing a minimum spanning network, not considering frequencies of haplotypes. This major clade with 21 haplotypes had good statistical support (89%), covering a large geographical range (Fig. 2b,c). Within the light purple clade, a total of 6 haplotypes or sub-clades was supported with good statistical bootstrap values between 77-93%.

In figure 2b the phylogeny of the 13 haplotypes in Lake Tinnsjøen revealed moderate to high bootstrap support for clustering of three “clades”; clade I (*h1, h2, h10*) with bootstrap support of 88%, clade II (*h5-h9. h11, h12*) with bootstrap support of 93%, and clade III (*h3, h4*) with bootstrap support of 85%. Here, clade I consisted of more haplotypes (i.e. *h13-18, h21, h32, h33*) that were found outside Lake Tinnsjøen and the four Norwegian outgroup lakes. One haplotype link, *h5*-*h13*, had unresolved cluster groupings, where it was interpreted that *h5*, being one mutational step away from *h1*, belonged to clade II rather than to clade I, and that *h13* belonged to clade I. The tree topology in figure 2b support these evaluations.

The minimum spanning network drawn using only the 13 haplotypes in Lake Tinnsjøen revealed variable frequency and their internal phylogenetic relationship (Fig. 2d).

Using the FA-morphs within Lake Tinnsjøen as units, the number of haplotypes ranged from 4 in the piscivore morph to 6 in the dwarf and planktivore morph (Table 3).

The percentage occurrence (based on Table 3) of the three clades in the four FA-morphs showed that the planktivore morph consisted of mostly clade III (77.3%), and less of clade II (18.2%) and clade I (5%). The dwarf had most of clade I (47.6%) and clade II (38.1%), and less of clade III (14.3%). The piscivore morph had mostly clade III (66.7%), and less of clade I (28.6%) and clade II (4.8%). Finally, the abyssal morph had most of clade II (72.7%), and less of clade III (18.2%) and clade I (9.1%).

The genetic diversity (Table 3) of FA-morphs ranged from a low haplotype diversity of 0.476 (planktivore morph) to a high 0.743 (dwarf morph) in Lake Tinnsjøen, and from 0-0.222 (highest in Lake Vatnevatnet) in outgroup lakes. In Lake Tinnsjøen combined, the haplotype diversity was 0.711. Similarly for nucleotide diversity, a low value was observed for the abyssal morph (0.00078) and a high value for the dwarf morph (0.00128), while the four outgroup lakes varied 0-0.00026 (highest in Lake Vatnevatnet). In Lake Tinnsjøen combined, nucleotide diversity was 0.00124.

Pairwise distance F_ST_ values based mtDNA Cytochrome B among the four morphs in Lake Tinnsjøen ranged from a low 0.042 (planktivore vs piscivore morphs) to a high 0.38 (planktivore vs dwarf) (Additional file: Table S4). All other F_ST_ comparisons than the planktivore versus the piscivore morphs were significantly different.

With regard to signals of demographic changes in mtDNA, support for demographic expansion was indicated for clade I (Tajimàs D; p-value > 0.05 and Fu and Lìs D; p-value 0.05) which comprised all four Norwegian outgroup lakes and 19 fish from Lake Tinsjøen (all four morphs present) (Additional file: Table S5). A stronger support for demographic expansion was suggested for clade II (Tajimàs D; p-value < 0.05 and Fu and Lìs D; p-value <0.02) which were endemic in Lake Tinnsjøen. However, no support for population expansion was suggested for the other endemic clade III (Tajimàs D; p-value > 0.10 and Fu and Lìs D; p-value > 0.10).

### Reproductive isolation of field assigned morphs or fish assessed using unbiased methods

The hierarchical STRUCTURE analysis suggested K=8 genetic clusters with the four morphs in Lake Tinnsjøen and the four Norwegian outgroup lakes occurred as distinct clusters (Fig. 5ad, Additional file: Table S9, hierarchical STRUCTURE plot in Additional file: Fig. S1).

The number of alleles in the four morphs and the four outgroup lakes ranged between 76 (Lake Vatnevatn) to 143 (planktivore morph), private allele richness from 0.13 (piscivore) to 0.69 (River Leirfossvassdraget), allelic richness from 6.02 (Lake Vatnevatn) to 8.63 (planktivore morph), F_is_ from -0.012 (Lake Tyrivann and Femund) to 0.118 (River Leirfossvassdraget), heterozygosity from 0.128 (piscivore morph) to 0.820 (Lake Tyrivann and Femund), and gene diversity from 0.567 (Lake Vatnevatn) to 0.761 (River Leirfossvassdraget).

Microsatellite based F_ST_ of morphs and sample lakes ranged from 0.08-0.29 (Additional file: Table S10). When using Lake Tinnsjøen as one group compared with the four Norwegian outgroup lakes, F_ST_ ranged 0.06-0.27 (Additional file: Table S11). If considering genetically pure GA morphs only (i.e. q>0.7), F_ST_ ranged between 0.09-0.21 (Additional file: Table 12).

On a broader scale, using the principal component analysis on microsatellite data (q=2.74, P=0.05), comparing the five lakes, revealed that all lakes were significantly different along PC1 and PC2 (Steel-Dwass method; q=2.72, alpha=0.05) except Lake Tyrivann and Lake Femund that were not significantly differentiated (Fig. 5b).

**Fig. 5.**
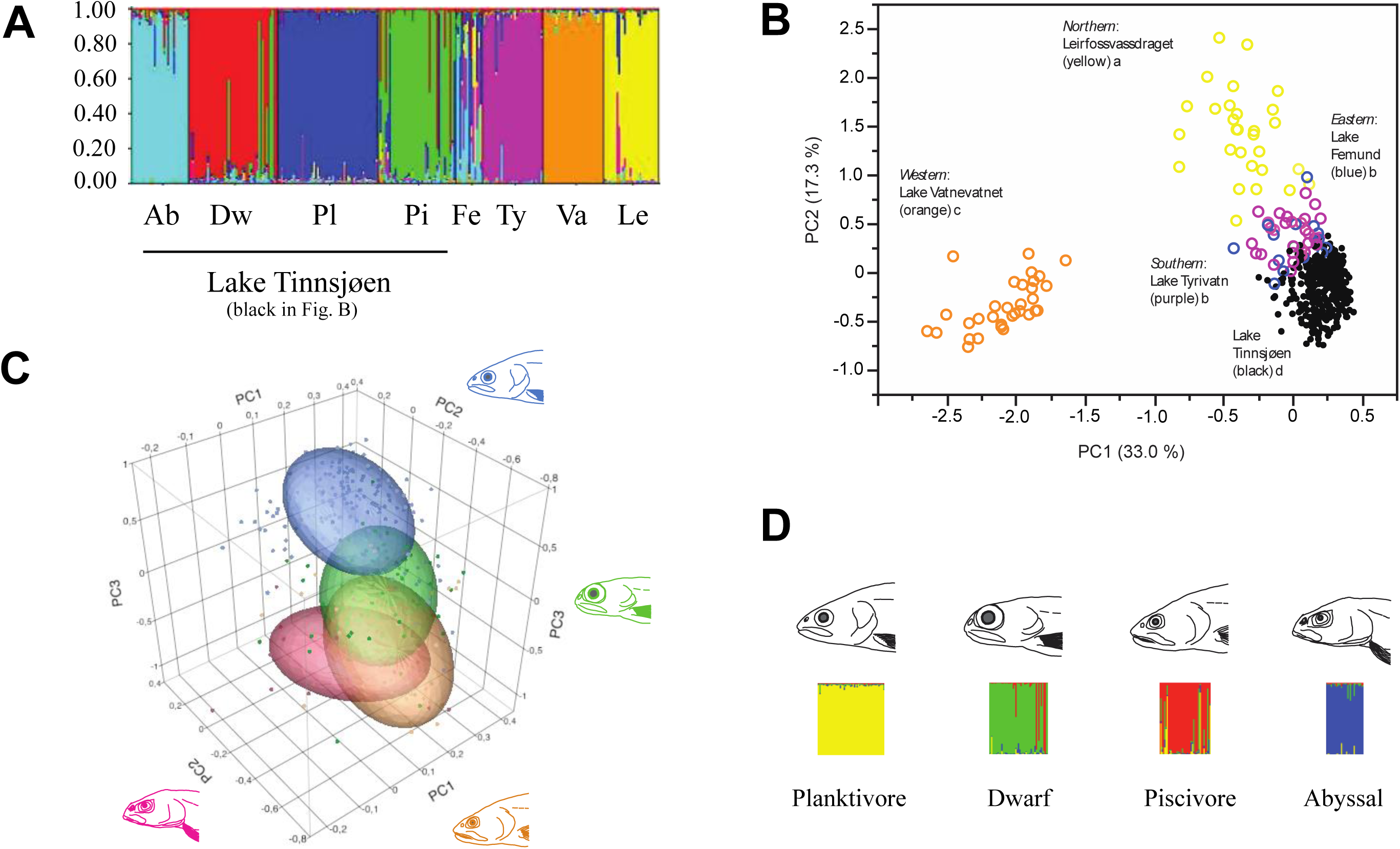
**(A)** STRUCTURE plot for K=8 genetic clusters based on the 10 microsatellites for the four Lake Tinnsjøen FA-morphs and for the four Norwegian outgroup lakes. Abbrevations; Lake Tinnsjøen (Ab=abyssal morph, Dw=dwarf morph, Pl=planktivore morph, Pi=piscivore morph), Fe=Lake Femund, Ty=Lake Tyrivatn, Va=Lake Vatnevatnet and Le=Leirfoss-vassdraget River. **(B)** PCA plot of microsatellite alleles partitioned into the five lakes studied (different letters denote significant differences on PC1; colors match figure A). (**C**) Three-dimensional PCA plot of microsatellite alleles for the four FA-morphs in Lake Tinnsjøen only (a subset of the four lakes visualized in figure B). The colours in graphs represents heads of the four FA-morphs given on the sides of the graph. (**D**) STRUCTURE plot for K=4 based on microsatellites in the four FA-morphs in Lake Tinnsjøen. Note that the colors in figure C and D are different and do not correspond to the same morphs across figures.

Using the same principal components on microsatellite as above, but only contrasting the four FA-morphs in Lake Tinnsjøen, revealed that four out of the six comparisons were significantly different for PC1 (q=2.57, alpha =0.05), and five of six were significantly different for PC2 (Fig. 5c). For PC1, the piscivore morph was not different from the abyssal morph, and the planktivore morph was not different from the dwarf. Along PC2, the dwarf morph was not different from the abyssal morph, while for PC3, the piscivore and abyssal morph did not differ significantly.

### Eco-morphological and life history trait divergence in the Lake Tinnsjøen charr morphs

In the contingency analysis of FA-morphs by habitat-specific catch the association was significant (N=457, Df=9, R^2^ (U)=0.400, Likelihood ratio test; χ^2^=387.92 and P<0.0001)(Fig. 3b). The planktivore morph was caught in the pelagial (22.3% of the catch within morph), littoral (36.2%) and shallow-moderate profundal (41.5%), but not in the deep profundal (0%). The dwarf morph was primarily caught in the shallow-moderate profundal (98.8%) appearing at 20-70 m depths, and only rarely in the littoral (1.2%). The piscivore morph was primarily caught in the shallow-moderate profundal (95.2%), and rarely in the littoral (3.2%), and deep profundal (1.6%). The abyssal morph was only caught in the deep-profundal habitat (100.0%).

In the contingency analysis of habitat-specific catch by the four revealed GA-morphs the association was also significant (N=344, Df=12, R^2^ (U)=0.4283, Likelihood ratio test; χ^2^=302.55 and P<0.0001), although less than 20% of cells in the tests had expected count <5 (suggesting x^2^ to be suspect)(Additional file: Table S6). The same general pattern emerged as previously described for FA-morphs above in the FA-morphs by habitat-specific catch contingency analysis.

When testing concordance of body shape and FA-morphs in Lake Tinnsjøen it was a moderate-strong concordant assignment ranging between 54.8% (piscivore morph) and 83.0% (abyssal morph) (Table 4, Fig. 4a).

**Table 4.** Assignment percentage based on discriminant analysis of PC-axis 1-5 for body shape comparing the four FA-morphs within Lake Tinnsjøen.

| Unit compared | Individuals | Planktivore | Dwarf | Piscivore | Abyssal |
| --- | --- | --- | --- | --- | --- |
| Planktivore | 282 | **83.0** | 9.9 | 1.1 | 0.4 |
| Dwarf | 81 | 8.6 | **70.4** | 14.8 | 1.2 |
| Piscivore | 62 | - | 25.8 | **54.8** | 8.1 |
| Abyssal | 32 | - | - | 9.4 | **71.9** |

In the contingency analysis of FA-morphs and GA-morphs association was significant (N=344, Df=12, R^2^ (U)=0.563, Likelihood ratio test; χ^2^=453.75 and P<0.0001) (Table 5). Here, association ranged from moderate 55.4% (dwarf morph) to a value of 100% (abyssal morph).

**Table 5.** Association between genetically assigned morphs (GA-morphs) based on microsatellite based STRUCTURE analysis (q>0.70) and the subjectively field-assigned morphs (FA-morphs). The group GA – hybrids is fish with a q-value < 0.70 and in such could not be assigned to any specific GA-morph. Values are percentages within morphs using genetic assignment in GA-morphs compared to FA-morphs.

| Unit compared | Individuals | FA- Planktivore | FA- Dwarf | FA- Piscivore | FA- Abyssal |
| --- | --- | --- | --- | --- | --- |
| GA - Planktivore | 166 | **94.6** | 3.0 | 2.4 | - |
| GA - Dwarf | 74 | 28.4 | **55.4** | 16.2 | - |
| GA - Piscivore | 41 | 4.9 | 17.9 | **78.0** | - |
| GA - Abyssal | 29 | - | - | - | **100.0** |
| GA - hybrids | 34 | 14.7 | 55.9 | 26.5 | 2.9 |

Similarly, back assignment using body shape of the FA-morphs and the putative ancestor from Lake Tyrivatn showed that Lake Tyrivatn had highest assignment to plantivore morph (18.8%), a lower assignment to dwarf (6.3%) and piscivore (3.1), while no fish from Lake Tyrivatn was assigned to the abyssal morph in Lake Tinnsjøen (Additional file: Table S7).

For comparative purposes, the FA-morphs and GA-morphs were visually contrasted with regard to age and weight distribution, suggesting large difference among morphs (Fig. 4bc). It seems that the planktivore morph has the lowest age span followed by a roughly equal life span in the dwarf and abyssal morph. The piscivore morph has the longest life span. There were large differences in weight distribution, where the piscivore attained the largest size followed by the planktivore morph. The dwarf and abyssal morphs were minute in comparison, but the dwarf morph attained a larger body size than the abyssal morph. The comparison of FA-morphs and GA-morphs broadly gave the same picture in age and weight.

## Discussion

In our study we have found empirical support for evaluating the three main research questions addressed. First, we find it reasonable to postulate that members of one Holarctically widespread mtDNA lineage colonized Lake Tinnsjøen, likely suggesting one single common ancestor that later diversified into the observed four sympatric morphs. Further, the number of endemic haplotypes, and signatures of population demographic expansion, in one of the Lake Tinnsjøen clades (clade II), support a mechanism of intralacustrine diversification. Secondly, we found that the four field assigned morphs were genetically divergent at microsatellite loci (and partly mtDNA), thus suggesting reproductive isolation among morphs (although with some degree of gene flow). Further, there were a close association between field assigned morphs and unbiased genetic analyses (microsatellites) revealing four distinct genetic clusters in the lake, supporting morph differentiation. Thirdly, we evaluate that the four morphs were differentiated with regard to habitat use based on catch, and in their life history, suggesting diversification along a depth-temperature-productivity-pressure gradient. Given that this adaptive radiation occurred after the lake became ice-free (<10 000 years), it represents a rapid diversification in lake niches with associated phenotypical modifications. If considering a 5 year mean generation time it correspond to a maximum of 2000 generations of evolution. Although not directly studied, the seen association between phenotypic divergence and catch habitat imply adaptive niche proliferation with morphological specialization (regardless of phenotypic plasticity or genomic hardwiring) towards different environmental conditions along the depth-temperature-productivity-pressure gradient in the lake. The degree of genetic differentiation is complementary to the level seen in other sympatric Arctic char lake systems. However, the degree of morphological differentiation, and niche radiation, in Lake Tinnsjøen reveal an extension of specialization into the deep profundal niche. Thus, this highlight an intriguing general question in speciation research of polymorphic fish in lakes; have we systematically underestimated the degree and rate of adaptive radiation into profundal niches?

### Timeframe of fish colonization into Lake Tinnsjøen based on the glacial geology pattern

It is relevant to address the glacial geological conditions surrounding the area of Lake Tinnsjøen for evaluating the potential of colonization direction and timing of founder events. The maximum extension of the Eurasian Late Weichelian ice sheet occurred ca 21-23 000 years before present (ybp) [36, 37]. Around 15 000 ybp the retreating ice margin was close to the Norwegian coast, and the ice stream in the Skagerak Sea broke up in the Norwegian channel [104]. In southern Telemark county, wherein Lake Tinnsjøen is situated, the ice sheet extended all the way to the coast ca 13 000 ybp [105]. Around 12 000 ybp the coast was ice free [104]. The ice-sheet retreated in a northwestern direction. An ice-recession line southeast of Lake Heddalsvatnet, situated below Lake Tinnsjøen in the same drainage (River Tinne), was dated to 9 700 ybp by Bergstrøm [105]. Further, marine sediment deposits was recorded (www.ngu.no) close to the village of Årlifoss 11 km southeast of Lake Tinnsjøen in River Tinne (see Fig. 1b for position of the upper limit of marine deposits). A sediment core study from Lake Skogstjern in the lower part of the Skiensvassdragets River by Wieckowska-Lüth et al. [106] revealed a lake formation dating at ca 10 500 ybp. The outlet of Lake Tinnsjøen is situated 50 km (estimated current waterway distance) northwest of Lake Heddalsvannet. Lake Tinnsjøen was glaciated and we thus assume that it could not have been accessible for fish immigration prior to that period – setting a crude frame for colonization to <9 700 ybp. We further infer that the fish colonization have proceeded from the southeast through the River Skienselva, or alternatively through any existing non-identified pro-glacial lakes situated southeast of Lake Tinnsjøen. This is also logic given the elevation level of the landscape surrounding Lake Tinnsjøen, where colonization along the suggested direction is most likely as the alternative routes imply crossing mountains and elevated slopes. Furthermore, the estimated ice-flow directions (Fig. 1b; [105],) support that the Arctic charr colonized Lake Tinnsjøen along the River Skienselva from the coastline and upwards. As the Arctic charr can be anadromous and live short periods in the sea [19], and as the Skagerak area at certain times during deglaciation was carrying a brackish water upper layer [107, 108], it seems reasonable to infer that the Arctic charr came from the south and colonized lake Tinnsjøen from the coast.

### Holarctic phylogenetic patterns using mtDNA CytB in Arctic charr and Salvelinus spp

We have screened a moderate number of each of the four morphs in Lake Tinnsjøen and only few fish in the four comparative Norwegian populations to the South, West, East and North of Lake Tinnsjøen for one mtDNA Cytochrome B fragment. In our evaluation, all these lakes hold Arctic charr as we would phenotypically recognize them in Norway, with perhaps the exception of the abyssal morph which is morphologically distinct. We used our sequences to obtain similar sequences from 18 named species of *Salvelinus* from GenBank to assess Holarctic patterns in haplotypes, lineages, clades and taxa distributions (Additional file: Table S2a). The initial description of taxa in GenBank sequences comprised; *Salvelinus* spp, *Salvelinus albus*, *S. andrashevi*, *S. alpinus*, *S. alpinus alpinus*, *S. alpinus erythrinus*, *S. boganidae*, *S. confluentus*, *S. elgyticus*, *S. krogiusae*, *S. kronocius*, *S. kuznetzovi*, *S. malma*, *S. malma malma*, *S. malma lordi*, *S. neiva*, *S. oquassa*, *S. schmidti*, and *S. taranetzi*. Here, we used the sequences from GenBank regardless of species denomination in the initial deposition or publications. This strategy was chosen as the species designation in the *Salvelinus* complex is challenging due to the traditional use of morphological species criteria with variable consensus. When considering all 88 haplotypes together, *S. alpinus* was described with haplotypes; *h1-h17*, and *h82*, while remaining taxa belonged to other taxa than *S. alpinus*. Here, Norway, Sweden, Finland, Canada, USA, and Russia report haplotypes of *S. alpinus*. Our Holarctic CytB phylogeny, and phylogeographic patterns, reveals a much deeper divergence than only considering the single Arctic charr *S. alpinus* taxa. As mtDNA is maternally inherited it only partly reveals the evolutionary history of the *S. alpinus* species complex. In general, one should be cautious inferring mtDNA based phylogeography as we assume selective neutrality, which may not always be the case [e.g. see 109].

For the five Norwegian lakes combined, we observed a set of 13 haplotypes (*h1-h13*). Haplotype *h1* was found in all the five Norwegian lakes, as well as in some other Holarctic lakes (Sweden, Finland, Russia, and Canada). Haplotype *h13* was found in Lake Tinnsjøen and one other lake in Sweden. Haplotype *h2* was only found in one Norwegian lake (Lake Vatnevatnet). A set of 10 haplotypes (*h3-h12*) were found to be endemic in Lake Tinnsjøen. In the phylogenetic analyses using the full dataset of the 88 mtDNA CytB sequences, a moderate to strong statistical support for branching events were observed (Fig. 3). These major branches were found in different parts of the Holarctic and reflects a deeper taxonomic partitioning than only containing Arctic charr (Additional file: Table S2a). However, our main purpose of this large scale comparison of CytB sequences (using different described charr (*Salvelinus* spp.) taxa) was to visualize, and polarize, the closest relatives in the major branch that also contained the Arctic charr in Lake Tinnsjøen. With that in mind, we focused on the major light purple lineage in Figure 3 (named #1 in Fig. 3a,b). This lineage included a set of 21 haplotypes widely distributed (Fig. 3c) in Transbaikalia - Kamchatka - Bering Sea (*h17*, *h18*, *h21*, *h32*, *h33*), Quebec - Taimyr - Chucotka - Fennoscandia (*h1*, *h2*, *h13, h14*, *h15*, *h16*), and in Lake Tinnsjøen (*h3*-*h13*). Based on these results, it seems plausible to evaluate that the ancestors of this major lineage colonized a large geographic area throughout the Holarctic, where some ancestral individuals also colonized Lake Tinnsjøen. Here, the founders of Lake Tinnsjøen could potentiall have carried the *h1* haplotype (clade I), subsequently giving rise to clade II (*h5*, *h7*, *h8*, *h9*, *h11*, *h12*) and clade III (*h3*, *h4*) (Fig. 3d).

In our study, considering the major lineage 1 (light purple; named #1 in in Fig. 3a,b) with 21 haplotypes (*h1*-*h18*, *h21*, *h32*, *h33*), it is interesting to see what species designations are given to haplotypes as these should be the closest relatives to morphs in Lake Tinnsjøen. This lineage was distributed in Quebec - Bering Sea - Kamchatka - Chucotka - Transbaikalia - Taimyr - Fennoscandia, and Lake Tinnsjøen. The taxa described were: *S. alpinus* (*h1-h13*), *S. a. alpinus* (*h14-h15), S. alpinus* and *S. a. oquassa* (*h16), S. alpinus*, *S. boganidae* and *S. a. erythrinus* (*h17), S. malma and S. m. malma (h18), S. malma (h21),* and *S. a. erythrinus* (*h32, h 33)*. Based on Osinov et al. [110], which used mtDNA CytB to assess phylogenetic relationship among taxa in *Salvelinus*, it seems that *S. alpinus*, *S. a. oquassa, S. a. erythrinus and S. malma* group together (MP tree), being genetically related to *S. boganidae* (evaluated from lack of bootstrap support). Thus, our results seems concordant with Osinov et al. [2015], and other studies regarding a genetic relationship between *S. malma* and *S. alpinus* [111–114], comprising taxonomic members in lineage 1 (light purple; named #1 in in Fig. 3a,b).

In general, there were few similar CytB sequences in GenBank, and specifically with regard to the Fennoscandian area (as revealed in Fig. 3d). However, we contrast our findings with the large scale phylogeographical study by Brunner et al. [115] who targeted Holarctic *Salvelinus* spp. using the mtDNA control region. They found five major phylogegraphic lineages, named; Atlantic, Acadia, Siberia, Bering, and Arctic groups in the *Salvelinus sp.* complex. Our phylogeographic data has much less geographical coverage than Brunner et al. [115], however, with some putative similarities. As such, our lineage 2 (yellow; named #2 in in Fig. 3a,b) may potentially fit with their Bering lineage, our lineage 5 (green; named #5 in in Fig. 3a,b) may fit with their Arctic lineage, and our lineage 1 (light purple; named #1 in in Fig. 3a,b) may fit with their Atlantic lineage. However, our lineage 1 seems to extend further north and east than the Atlantic lineage revealed by Brunner et al. [115]. A study by Gordeeva et al. [116] studying Arctic charr (mtDNA control region) in the European part of Russia and Siberia revealed that the Atlantic group was distributed all the way to Taimyr (see also [117]). Further, Alekseyev et al. [118] found no strong support for a separation of Atlantic and Siberian haplotypes into two distinctive groups when analyzing Arctic charr in Siberia using the mtDNA control region. Thus, these studies in general seem to support our crude inference of a widespread Atlantic mtDNA lineage (Fig. 3a,b). However, we seem to lack the Siberian, the Acadian, and the Svet lineage as being revealed by Brunner et al. [115]. The discrepancy between our results and Brunner et al. [115] may be due to lower geographical coverage, use of CytB as a much less powerful marker for divergence than the control region, and also a different set of individuals that may, or may not, belong to different *Salvelinus* taxa proposed.

### Patterns in adaptive radiation of Arctic charr – genetic divergence of sympatric morphs

In the Holarctic it seems to be a pattern in adaptive diversification into lake niches in Arctic charr where most lakes hold only one morph (e.g. littoral), fewer lakes have two morphs (e.g. littoral and pelagic), even fewer lakes have three morphs (e.g. littoral-pelagic and profundal), while only one lake so far has been reported to harbor four morphs (small and large benthic, planktivore and piscivore) (see relevant references below). Here, we contrast lakes with Arctic charr holding one, two, three or four morphs with regard to genetic differentiation in microsatellites unless otherwise stated. The comparisons and case systems are not complete, meaning that we do not cover all systems, but assumes that this is a representative set of polymorphic systems in the Holarctic. The measure for evaluating genetic differentiation is the common metric F_ST_ if not stated otherwise. The different microsatellite loci applied and significance levels of F_ST_ value comparisons among morphs are found in the given references.

Several studies have compared Arctic charr among lakes with regard to their genetic differentiation (where there may be lakes holding more than one morph of Arctic charr) revealing a F_ST_ range of 0.003-0.627 when contrasted in Holarctic lakes [119–127]. Presence of two morphs associated (or not) with genetic clusters have been found in a number of Arctic charr lakes revealing a F_ST_ range of 0.001-0.381 in Holarctic lakes [55, 60, 62, 66, 123–125, 128–133]. When considering Arctic charr lakes with support for three morphs and/or genetic clusters, much fewer systems are reported. Moccetti et al. [62] report three morphs (littoral omnivorous, small-sized profundal benthivorous, and large-sized profundal piscivorous) in Lake Tårnvatn in Norway with a range in F_ST_ 0.042-0.134. Gíslason et al. [128] compared three morphs (planktivore, piscivore and benthivore) in Iceland in Lake Svinavatn with F_ST_ of 0-0.059. A later study by Wilson et al. [123] estimating F_ST_ 0-0.085; suggesting only two morphs in that particular lake. May-McNally et al. [126] studied three Alaskan Arctic charr morphs (large, medium and small-bodied) in Lake Tazimina with F_ST_ 0.017-0.092. Alekseyev et al. [134] studied three Arctic charr morphs (dwarf benthophage, small planktophage and large predator) in Lake Kamkanda in Transbaikalia and found F_ST_ 0.168-0.299. Gordeeva et al. [60] studied three lakes with three morphs in Transbaikalia, Russia revealing F_ST_ 0.015-0.497. Across Holarctic lakes with three morphs, a range in F_ST_ values from 0 to 0.497 were observed. A set of four morphs (small and large dark and small and large pale morphs) have been described from Gander Lake in Canada [135, 136]. Gomez-Uchida et al. [137] tested the dark and pale morphs and found F_ST_ (theta) 0.136, suggesting two genetic clusters. Currently, it is unknown whether these four morphs constitute four genetic clusters. The classic textbook example of adaptive radiation in Arctic charr comes from a continental plate rift lava lake in Iceland. Here, a set of four morphs of Arctic charr are described in Lake Thingvallavatn; large benthic, small benthic, planktivorous and piscivorous morphs [59]. Kapreolova et al. [125] studied three of these morphs (small benthic, large benthic, planktivorous,) and found F_ST_ (theta) 0-0.07. As such, the genetic status of the four Lake Thingvallavatn morphs remains partly unresolved to date with regard to microsatellite differentiation. In our study of the Arctic charr in Lake Tinnsjøen, we estimated F_ST_ 0.130-0.195 among the four morphs, being much more differentiated than among the three compared morphs in Lake Thingvallavatn. However, the range in genetic differentiation among morphs in Lake Tinnsjøen lies within the range among-lakes (F_ST_ 0.060-0.627), among two-morph sympatric systems (F_ST_ 0.010-0.381), and within the three-morph sympatric systems (F_ST_ 0-0.497). It is difficult to discuss in more detail given the large F_ST_ range in the four types of lakes (mono-quadrets). Also, the different marker sets used in various studies makes it difficult to conduct a direct comparison, but the level of genetic segregation within Lake Tinnsjøen seems generally to comply with the level of genetic segregation seen in other sympatric systems. As Lake Tinnsjøen is the only lake with four morphs of Arctic charr, currently known, the range in F_ST_ was 0.130-0.195.

With regard to mtDNA divergence (and other marker sets) of sympatric Arctic charr morphs much fewer studies exist. Alekseyev et al. [118] studied a set of 22 lakes in Transbaikalia, Russia, analyzing the mtDNA control region. They found that sympatric morphs shared one or two haplotypes in each lake, with no significant differentiation in haplotypes between morphs in each of the lakes. Salisbury et al. [138] found that all their Arctic charr in Gander Lake belonged to the Atlantic mtDNA D-loop lineage (except one pale morph individual). They implied that morphological, ecological and genetic differentiation of dark and pale morphs could be due to sympatric origin within the last 10 000 year postglacial timeframe. Alekseyev et al. [118] and Gordeeva et al. [60] studied the same 22 lakes having sympatric morphs in Transbaikalia, Russia. They suggested that six of these lakes may represent evolutionary events of independent parallel divergence in sympatry, since they had their own set of endemic mtDNA-control region haplotypes shared only among morphs within lakes, while 16 lakes could could evaluated as potential allopatric events due to sharing of mtDNA haplotypes both among morphs and nearby lakes. Three morphs of Arctic charr were suggested in Loch Rannoch in Scotland by Verspoor et al. [139] being the pelagic morph and two benthic morphs (small and large mouth). They analysed mtDNA-RFLP in the D-loop and found that the pelagic morph was divergent from the benthic morphs, with F_ST_ between the pelagic and the two benthic morphs ranging 0.326-0.487. Differentiation among the two benthic morphs was lower with F_ST_ 0.158. The authors suggested that the pelagic morph and the two morphs comprised two allopatric lineages, while a sympatric divergence could have led to the two benthic morphs after colonization of their ancestral mtDNA lineage into the lake. In the four morph system in Lake Thingvallavatn, Magnusson and Ferguson [140] analyzed allozymes and found that all four morphs were genetically closely related with a Neís (D) range 0.00004-0.00126, suggesting that the four morphs did not belong to different evolutionary lineages. They stated that caution should be used interpreting data using few polymorphic loci. Later, Volpe and Ferguson [141] analysed sequences and mtDNA restriction fragment analysis of the control region, and minisatellites, but according to the authors own statement lacked resolution to test the specific hypotheses regarding sympatric origin and genetic divergence among morphs. However, they found some support for morphs in the lake to be more genetically similar than to other morphs in other lakes, potentially supporting an intra-lacustrine diversification (although low bootstrap support for divergence of the morphs within Lake Thingvallavatn). Danzmann et al. [142] used mtDNA restriction analyses and suggested that the Thingvallavatn morphs were closely related. Escudero [143] analysed the d-loop in three of the morphs (planktivorous, small and large benthic morphs; not including the piscivore morph) and found five haplotypes where two haplotypes were shared among all morphs, one morph had two endemic haplotypes, and another morph had one endemic haplotype. Barring a full contrast of the four morphs with regard to mtDNA, there seems to be no strong differentiation in mtDNA among morphs in this lake. However, Lake Thingvallavatn is indeed one of the best Arctic charr systems studied worldwide and two studies by Gudbrandsson et al. [144, 145] reveal presence of significant development transcriptomic gene expression differences among the four sympatric morphs, suggesting extensive genetic divergence among sympatric morphs. Gudbrandsson et al. [144, 145] also discuss an alternative hypothesis of whether or not the piscivore morph exists as one genetic cluster, if it has recently diverged under asymmetric gene flow from other morphs, or if it is an inducible morph due to threshold values in growth before becoming a piscivore. Based on these two studies, it seems that the piscivore morph is less genetically distinct than the three other morphs in that lake. In comparison, the F_ST_ values among the four sympatric morph in Lake Tinnsjøen were 0.042-0.382. Here, the only comparison that was non-significant was between the planktivore and the piscivore morph. Thus, five out of the conducted six morph comparisons in Lake Tinnsjøen were significantly different, implying limited gene flow among morphs. The number of haplotypes ranged from 4-6 among the four morphs, with the number of endemic haplotypes differing from 1-2 among morphs. There is a possibility that the small sample size in our mtDNA analyses reflect biased sampling of morphs, but that do not seem likely given the sample size of 21-22 individuals from each morph. Due to lack of previous studies, it appears that no direct comparison can be made to relevant studies on Arctic charr with regard to contrasting F_ST_ values based on mtDNA. However, using the same line of argument as in Alekseyev et al. [118] and Gordeeva et al. [116], one could imply a case of sympatric origin of the four lake Tinnsjøen morphs as they have endemic haplotypes not yet seen outside the lake. However, that could also reflect limited geographical coverage nearby, or far from, Lake Tinnsjøen. Thus, one should be cautious interpreting these results.

In summary, genetic divergence (using different markers) among sympatric Arctic charr morphs in lakes throughout the Holarctic varies widely, and we expect them to do so given their different evolutionary histories, genetic load and evolvability, biotic and abiotic environmental conditions, and ecological opportunities to radiate. Indeed, we can see systems with one to four morphs in different lakes. However, few studies have addressed nuclear and mtDNA markers at the same time. The best studied system so far is Lake Thingvallavatn in Iceland, where four morphs reside – where one of the morphs (i.e. piscivore morph) could be recently evolved as a separate genetic cluster or being induced due to growth threshold dynamics. In comparison, in Lake Tinnsjøen we have described four morphs that are different with regard to microsatellites and mostly with regard to mtDNA haplotypes. The Arctic charr morphs in Lake Tinnsjøen seem to be differentially distributed along a depth-temperature-productivity-pressure gradient. The evolutionary branching in their phylogeny, and the high number of endemic haplotypes in Lake Tinnsjøen, with signatures of demographic expansion, could support an intra-lacustrine origin of these morphs. However, the evolutionary scenarios remain to be tested in detail using a set of higher resolution markers. Although the Arctic charr species complex has been studied for a long time, researchers still need to address the important mechanisms underlying origin, presence and temporal persistence of sympatric morphs. Thus, a multi-method based eco-evo-devo approach with ecological, morphological and life history studies [146], and state of the art genomics as performed in Lake Tingvallavatn [e.g. 144, 145], seem to be a good avenue, as well as the methods applied in Jacobs et al. [23] contrasting two independent replicate lineage radiations of the Arctic charr.

### Adaptive divergence of morphs - yes likely - but, what are the drivers of diversification?

Is there a repeatable pattern in niche use in sympatric morph? Imagine the colonization of a barren lake after the ice age with all lake niche available for utilization. Here, founders will likely utilize the most energetically profitable niche first, depending upon the lake-specific morphometry with regard to the highest fitness gain in the littoral or pelagial niche. Thus, the starting point for adaptive proliferation may be highly contingent on what niche(s) is actually holding the highest fitness reward among the available lake niches. This will also apply in a situation with presence of another species being a resource competitor or predator. Imagine a shallow lake with large littoral zone and a small pelagic zone. Here, it is reasonable to expect a higher temporally reliable food production in the littoral producing a higher proportion of littoral adapted morph individuals and a smaller proportion of pelagial adapted morph individuals. If the lifetime fitness is higher in the littoral than in the pelagial, radiation may not occur, but be present as a temporal utilization of the pelagic zone during zooplankton blooms in summer. As such, the pelagial may not harbour available energetic capacity during the whole year to open for a permanent life style adapted to strictly the pelagial. This could be an explanation for the many Arctic charr lakes that seem to hold monomorphic populations, acting as generalists. In contrast, deeper lakes, or a deep fjord lake, where the pelagial is proportionally much larger than the littoral, the pelagial may have the highest food resource and the expected lifetime fitness reward. Here, one could imagine that most fish would adapt to the pelagial and less to the littoral. In a given lake, if both habitats hold temporal stable and predictable energetically rewarding resources, one may expect that two morphs can evolve, one in the pelagial and one in the littoral, likely with relative proportions of morphs associated with relative proportion of habitat available according to an ideal free distribution. The same would apply for a system with three niches and morphs – evolving a morph adapted to the profundal. Based on the number of sequence of morphs from monomorphic to four morph systems, it seems that there is a predictable temporal pattern in evolutionary branching associated with niche radiation. Here, the littoral (or pelagial) may be the first niche to be filled – then the pelagial (or littoral) – then the profundal, with a piscivore morph originating putatively due to growth threshold dynamics from one of the units, or evolving independently.

Adding upon this complexity, moving away from an assumption of only three discrete niches in given a lake, one can imagine that there could be gradients of expected predictable fitness along environmental variation such as e,g, the depth-temperature-productivity-pressure gradient in Lake Tinnsjøen. Indeed, a study on polymorphic European whitefish (*Coregonus lavaretus*) in the Swizz Alpine Lake Neuchâtel suggest adaptive diversification and build up of reproductive isolation along ecological gradients when assessing morphs spawning at different time and place [147]. Ohlberger et al. [148] used an adaptive-dynamics model, calibrated with empirical data, finding support for an evolutionary diversification of the two German Lake Stechelin *Coregonus* sp. morphs likely driven by selection for physiologically depth-related optimal temperatures. In the 1.6 km deep Lake Baikal, Russia, one of the oldest freshwater lakes on earth, adaptive radiations have occurred in several taxa such as e.g. reflected by the depth-gradient and the environmental niche radiation of the freshwater sculpins (*Cottidae*, *Abyssocottidae* and *Comephoridae*) [149]. Also, speciation along depth gradients in the ocean are strongly suggested [150]. A study by Chavarie et al. [151] tested a multi-trait depth gradient diversification of morphs in Lake trout (*Salvelinus namaycush*) in Bear Lake in Canada, but did not find a strong association in differentiation with depth (but, partly association with genetic structure), suggesting that a highly variable nature of ecological opportunities existed for divergent selection and phenotypic plasticity. In comparison with these studies, it seems reasonable to infer that there is a depth-temperature-productivity-pressure gradient with different fitness rewards reflecting an adaptive landscape where upon the four Arctic charr morphs within Lake Tinnsjøen can adapt. Such a gradient may not necessarily be discrete with regard to environmental sustainable conditions, but could reflect a continuum, or a holey adaptive landscape [see 26]. A recent study by Jacobs et al. [23] reveal the complexity in inferring mechanisms behind origin of replicate Arctic charr morphs. These authors suggested that similar morphs, contrasting the Atlantic and Siberian lineage of Arctic charr, could originate through parallel or non-parallel evolutionary routes as revealed in gene expression being highly similar between independently derived replicates of the same morph. They highlighted that variability in the Arctic charr with regard to predicting phenotypes was contingent on a set of factors being demographic history, selection response, environmental variation, genomic architecture and genetic association with specific morphs. Thus, revealing mechanisms in speciation trajectories in the Arctic charr complex is indeed a challenging task.

A novel finding in our study was the appearance of the deep-profundal abyssal morph with its distinctive phenotypic features, apparently being adaptations to the cold, dark and low-productive high-pressure environment in deeper parts of the oligotrophic Lake Tinnsjøen. Our finding of the four morphs could reflect ongoing divergence along a depth-temperature-productivity-pressure gradient from surface to deep profundal environments. This imply large differences in yearly cumulative temperature sum at different depths and productivity, likely strongly affecting life history evolution. In shallow Fennoscandian lakes, the littoral seem to have the highest biotic production, followed by the pelagial and profundal [152]. In the 1.6 km deep Lake Baikal, oligochaetes was found from the surface down to maximum depth, comprising up to 70-90% of biomass and numbers in the bottom fauna [153]. In the same lake, biomass of benthos decreased with depth, with an increasing proportion of oligochaetes. In comparison with the Baikal studies, we assume that the biotic prey production for Arctic charr is highest in the pelagial in the deep Lake Tinnsjøen (with small littoral areas) and lower in the benthic-littoral, and the least in the deep profundal. As such, a temperature and food production gradient likely exists in Lake Tinnsjøen from more productive pelagic and littoral areas down to the shallow profundal and deep profundal. Also, as the pressure increase by one atmosphere each 10 meters depth, it should also have marked impacts on adaptations evolved in various traits, being particularly evident in the small abyssal morph with its curved head, upturned mouth and small eye size. Thus, both abiotic factors, and ecological opportunity, likely determine potential of adaptive divergence in deep water lakes as already implied in studies on Arctic charr in the profundal habitat [19, 53]. In deep lakes such as Tinnsjøen (460 m) and Gander Lake in Canada (288 m; [154]) selective forces for habitat and niche occupation could be even stronger than previously anticipated, selecting traits that have not been seen in other morphs from other lakes. In Lake Tinnsjøen, the small eyes (an apparent reduction of size and potential function?) in the abyssal morph bear apparent similarities with eye-reduction seen in cave fishes [e.g. 155]. This seems somehow logical given that cave environments often can be described as nutrient-poor, cold, and harboring few present species.

In speculation, there might be a temporal cascade effect in the adaptation process to a given niche where e.g. body size, growth rate, sexual maturation, coloration, secondary sexual traits, physiology and morphology are highly contingent upon the ecological opportunity, constraining environmental conditions, and the evolutionary optimal solutions in any given niche. Here, e.g. morphology and physiology may reflect specialization to niches, growth rate and size and age at sexual maturity may reflect food conditions and predation regimes, while body size, coloration and secondary sexual traits may reflect the optimal visual conditions, affecting mate choice behavior and thus sexual selection. Here, evolution would likely result in optimal solutions to obtain the highest overall life-time fitness in a given niche, and also due to relative fitness rewards in yet other niches in the lake due to overall e.g. frequency or density dependent fitness of the present morphs. Further, in this adaptive process e.g. major histocompatability complex (MHC) genes, which are related to e.g. kin recognition, parasite and disease resistance, as well as niche occupation, may be important as previously shown to differ among morphs and lakes in Arctic charr [156–159]. A study by Baillie et al. [160] surveying microsatellite and a MHC gene in Lake Trout in Lake Superior revealed that variation was partitioned more by water stratum than by ecomorph with a stronger association with MHC gene variants and depth. This suggests presence of an adaptive MHC gene polymorphism along the depth gradient, potentially due to divergent environments and ecological niches. As Arctic charr has sexual dimorphism in coloration it also implies sexual selection as a driver for population or morph differentiation. The high level of genetic differentiation often seen in sympatric Arctic charr morphs in comparison with e.g. a lower inter-morph genetic differentiation in European whitefish radiations [e.g. 133, 161] may suggest that sexual selection may be less pronounced in whitefish than in Arctic charr. Coloration in Arctic charr may thus be a reliable evolutionary signal of parasite and disease resistance, as well as niche use, used in mate choice preferences. One should also in such studies consider the optimal color wavelengths present at different depths or niches.

It is pertinent to pose the question whether the Lake Tinnsjøen morphs have originated due to ecological speciation mechanisms. According to the ecological theory of adaptive radiation and ecological speciation [3, 16, 48, 162], our four morphs do seem to fit well to an ongoing diversification process according to several of the expectations from this theory (see also [163–167]). However, the process of ecological speciation is complex and remains to be tested awaiting ecological niche studies and using higher resolution genetic markers under an evolutionary scenario framework comparing simulated and empirical data. As a crucial and fundamental basis in ecological theory, we would also here, in our newly discovered Lake Tinnsjøen system, expect a niche-specific fitness trade-off in adaptations to evolve so that no one phenotype will be optimal in all the available lake niches. Thus, the saying “*Jack of all trades, master of none, but oftentimes better than master of one*” might nicely reflect the early postglacial stages of the ongoing evolutionary dynamics in adaptive radiation of Arctic charr.

## Conclusion

In Lake Tinnsjøen, we revealed four Arctic charr morphs associated with differential catch in four habitats in the pelagial (<20 m), littoral (<20 m), shallow - moderate profundal (20-150 m), and deep profundal (150-350m). Apparently, morphs diverge along a depth-temperature-productivity-pressure gradient. Field assignment from exterior appearance, and laboratory geometric landmark analyses, support distinction into four morphs. Life history parameters also supported morph separation based on size, age and maturity patterns. MtDNA implied colonization of founders from one widespread Holarctic lineage, with subsequent origin of two new clades in Lake Tinnsjøen. Most morph pairs were genetically differentiated for mtDNA (F_st_: 0.04-0.38). Microsatellites revealed significant reproductive segregation among all morph pairs (F_st_: 0.12-0.20), with presence of gene flow. A novel finding was the abyssal morph in the deep profundal which has not yet been described before in the worldwide Arctic charr species complex. Thus, the deep profundal needs to be studied more in polymorphic fish species complexes as we may have overlooked a substantial part of the present biodiversity below the species level. Whether or not Lake Tinnsjøen represents a true sympatric speciation process remains to be tested using a combined set of genetic markers to contrast evolutionary scenarios. Lake Tinnsjøen offers a rare research window into an ongoing speciation process – evidently revealing an important part of the worldwide evolutionary legacy of Arctic charr. We suggest that the Norwegian management authorities merit Lake Tinnsjøen special biodiversity protection as it is one of the most divergent Arctic charr systems seen worldwide.

## Declarations

### Ethics approval and consent to participate

Not applicable.

### Fishing license

Fish were sampled after initial consent from local authorities at Tinn County Administration giving us permission to fish in Lake Tinnsjøen after consent was approved also by the local landowners. The local landowners gave us the oral permission to fish on their land. No other permit or ethics approval is needed in Lake Tinnsjøen in order to sample Arctic charr when the main purpose is to use Arctic charr for scientific studies.

### Consent for publication

Not applicable.

### Availability of data and material

Data deposition: Parts of the data used in this study are available as online supplementary information in the electronic version of the article. The reason why not all data have been freely distributed is the current unknown status of the abyssal morph described in our survey. As such, conservation authorities should evaluate the taxonomic status and conservation need of the morph before information on specific sampling locations and catches may be released. This is a logic precautionary conservation biological approach as the observed new abyssal morph in the deep-profundal habitat (150-350 m) may have a small/vulnerable population size. Further, as this morph is not found elsewhere in the world, it merits the highest conservation status possible. The Lake Tinnsjøen represents a unique window into speciation for scientists.

### Authors’ contribution

K.P. and K.Ø. conceived/designed the study (equal project leaders). All authors contributed in the field sampling (with exception of A-M.P.T. who entered the project at a later stage). Basic genetic work in the laboratory was conducted by M.H.H, M.H. and K.P. Analyses of fish morphology was done by M.H.H. and M.H. Genetic analyses was done by K.P., M.H., A.-M.P.T. and K.Ø. The main body of the manuscript was written by K.Ø., K.P. and M.H. with significant contributions from all coauthors. All authors read and approved of final manuscript.

## Acknowledgements

We thank the local authorities at Tinn County Administration and all the landowners around Lake Tinnsjøen for permission to fish. A large number of people were involved in the field sampling which we highly appreciated; G. Sabalinkiene (RIP), A.-E.B. Stokkenes, J. Kosir and A.S. Josvanger. We also thank M. A. Svenning for allowing us to use the microsatellite data from River Leirfossvassdraget. A very special thanks go to *Tinn Jeger og Fiskeforening* (Tinn hunting and fishing organization) for loan of the nice boat and for all the help throughout the fish sampling! Thanks to E. Østbye for discussion and support in the project. Thanks to Olivier Devineau and Kimmo K. Kahilainen for discussions and comments that greatly improved the manuscript. Thanks to our funding institution Inland Norway university of Applied Sciences for economical support for the ongoing project and publication funding.

## ADDITIONAL FILE INFORMATION

**Additional file: Table S1.** Summary table for the different number of arctic char used in different analyses.

| Lake/source | FA-morph | Sampled<br>- N | Body<br>shape -<br>N | MtDNA<br>- N | Microsatellites<br>- N | Weigth<br>FA | Weigth<br>GA | Age<br>FA | Age<br>GA |
| --- | --- | --- | --- | --- | --- | --- | --- | --- | --- |
| Tinnsjøen | Planktivore | 282 | 266 | 22 | 185 | 282 | 166 | 85 | 55 |
|  | Dwarf | 81 | 78 | 20 | 73 | 81 | 74 | 34 | 30 |
|  | Piscivore | 62 | 55 | 22 | 57 | 62 | 41 | 37 | 35 |
|  | Abyssal | 32 | 32 | 22 | 30 | 32 | 29 | 26 | 25 |
|  | Hybrids | - | - | - | - | - | 34 | - | 10 |
| Tyrivann | - | 32 | 26 | 6 | 32 | - | - | - | - |
| Vatnevatn | - | 64 | - | 9 | 32 | - | - | - | - |
| Leirfoss | - | ? | - | 5 | 29 | - | - | - | - |
| Femund | - | 14 | - | 9 | 14 | - | - | - | - |
| Genbank | - | - | - | 77 | - | - | - | - | - |
| Sum |  | 567 | 457 | 192 | 452 | 457 | 344 | 182 | 155 |

**Additional file: Table S2a.** The 88 Cytochrome B - mtDNA haplotypes compared in Holarctic *Salvelinus* sp. including Lake Tinnsjøen and the four Norwegian outgroups and similar sequences in GenBank. This is a separate excel file not yet released.

**Additional file:Table S2b.** Description of 10 microsatellites that was used in arctic char in Lake Tinnsjøen and in four Norwegian outgroups.

| Locus | Range base pair (bp) in our study | Sequence repeat motif (see the reference in last column) | Initially described motif | GenBank accession number | Reference for microsatellite markers |
| --- | --- | --- | --- | --- | --- |
| OMM1105 <sup>1</sup> | 112-194 | (AGAC)<sub>23</sub> (GATA)<sub>16</sub> | Not defined | AF352768 | Rexroad et al. (2001) |
| SalF56SFU <sup>2</sup> | 183-211 | (TG)<sub>25</sub> | Dinucleotide | AF537307 | McGowan et al. (2004) |
| SalP61SFU <sup>3</sup> | 108-164 | (CA)<sub>18</sub> | Dinucleotide | AF537312 | McGowan et al. (2004) |
| Sco204 <sup>*</sup> | not given | (TCTA)<sub>29</sub> | Tetranucleotide | AY88873 | DeHaan and Arden (2005) |
| Sco212 <sup>4</sup> | 226-382 | (ATCT)<sub>18</sub> | Tetranucleotide | AY88881 | DeHaan and Arden (2005) |
| Sco218 <sup>5</sup> | 161-245 | (GATA)<sub>31</sub> | Tetranucleotide | AY88887 | DeHaan and Arden (2005) |
| Sco220 <sup>6</sup> | 239-367 | (ATAG)<sub>5</sub>(ATAC)<sub>2</sub>(ATAG)<sub>15</sub>- | Tetranucleotide | AY88889 | DeHaan and Arden (2005) |
| SMM17 <sup>7</sup> | 105-135 | (CA)<sub>29</sub> | Dinucleotide | AY327127 | Crane et al. (2004) |
| SMM22 <sup>8</sup> | 155-271 | (TAGA)<sub>19</sub> | Tetranucleotide | AY327129 | Crane et al. (2004) |
| SalJ81SFU <sup>9</sup> | 101-159 | (GT)<sub>33</sub> | Dinucleotide | AF537304 | McGowan et al. (2004) |
| Sco215 <sup>10</sup> | 275-283 | (GAAA)<sub>16</sub>(GA)<sub>6</sub>- | Tetranucleotide | AY88884 | DeHaan and Arden (2005) |

---

* This locus was not used in our analysis as it was in LD with another locus, thus no range in basepairs have been provided.

**Additional file: Table S2c.** The 10 microsatellites used in arctic char in Lake Tinnsjøen and the 4 Norwegian outgroups. This is a separate excel file not yet released.

**Additional file: Table S3.**
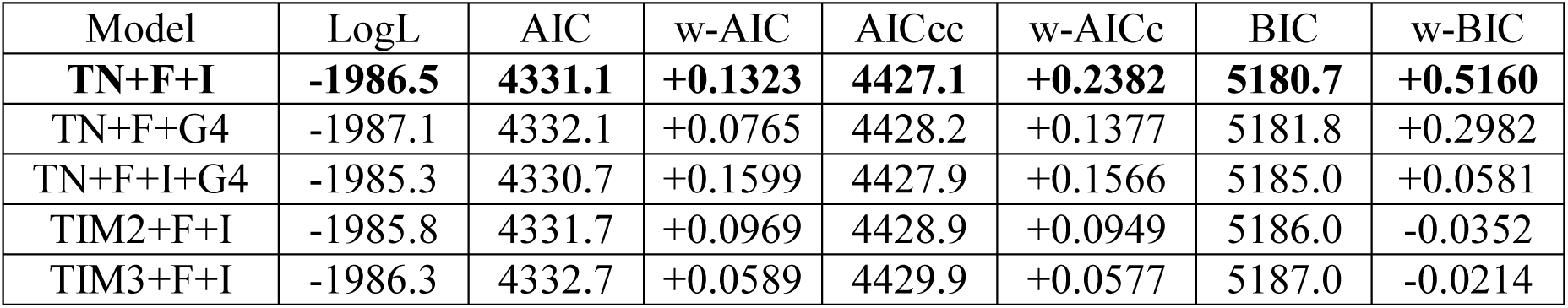
The best substitution model selection in IQ-Tree (http://www.iqtree.org/) [73–75; Nguyen et al. 2015, Kalyaanamoorthy et al. 2017, Hoang et al. 2018] with the best five models for the combined dataset of 88 haplotypes (13 Norwegian haplotypes and the 75 haplotypes found in Genbank). The best models selected was based on BIC in bold.

**Additional file: Table S4.**
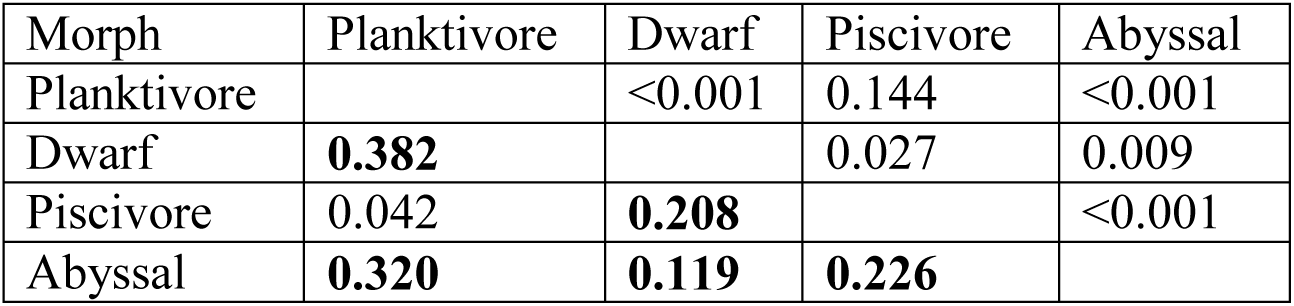
Pairwise distance mtDNA Cytochrome B F_ST_ values among the four FA-morphs in Lake Tinnsjøen. In the lower diagonal is given F_ST_ values while in the upper diagonal is presented p-values. Significant F_ST_ values are highlighted in bold.

**Additional file: Table S5.** Summary statistics for genetic-demographic analyses in DNAsp for the three clades that occur in Lake Tinnsjøen and the four Norwegian outgroup lakes. These clades have been selected based on the bootstrap support (>85%) (see Fig. 2b). Values in bold denote a significant test with regard to population expansion events.

| Parameter | Clade I | Clade II | Clade III |
| --- | --- | --- | --- |
| N basepairs | 851 | 851 | 851 |
| N sequences | 48 | 29 | 38 |
| N haplotypes | 4 | 7 | 2 |
| Variable sites/ singleton sites | 3/3 | 6/6 | 1/1 |
| Parsimony informative sites | 0 | 0 | 0 |
| Haplotype diversity | 0.122 | 0.377 | 0.053 |
| Nucleotide diversity (Pi) | 0.00015 | 0.00049 | 0.00006 |
| Mean pairwise differences k | 0.12500 | 0.41379 | 0.05263 |
| Theta per site from S, Theta-W | 0.00079 | 0.00180 | 0.00028 |
| Theta per sequence from S, Theta-W | 0.67599 | 1.52782 | 0.23801 |
| Raggedness r | 0.5882 | 0.1734 | 0.8033 |
| Ramos-Onsins & Rozas R<sup>2</sup> | 0.0807 | 0.0675 | 0.1601 |
| Theta initial | 0.000 | 0.000 | 0.000 |
| Tau | 0.125 | 0.414 | 0.053 |
| Tajima's D | -1.70014 | -2.09841 | -1.12863 |
| Tajima's D, P-value | > 0.05 | < **0.05** | > 0.10 |
| Fu and Li's D | -2.99741 | -3.39754 | -1.75813 |
| Fu and Li's D, P-value | < **0.05** | < **0.02** | > 0.10 |

**Additional file: Table S6.** Association between genetically assigned morphs (GA-morphs) based on STRUCTURE where “pure” morphs have q<0.7 and hybrids have q>0.7 and their catch in the four lake habitats. Values are percentages within morphs (rows) while the bottom row summarize overall percentage of catch in the four lake habitats. The last column summarize the overall percentage in the field catches with regard to relative percentage of GA-morphs.

| GA-morphs | Individuals | PEL | LIT | SDP | ABY | % of catch |
| --- | --- | --- | --- | --- | --- | --- |
| Planktivore | 166 | 19.3 | 33.1 | 47.6 | - | 48.3 |
| Dwarf | 74 | - | 5.4 | 93.2 | 1.4 | 21.5 |
| Piscivore | 41 | - | - | 100.0 | - | 11.9 |
| Abyssal | 29 | - | - | - | 100.0 | 8.4 |
| Hybrids | 34 | - | 2.9 | 94.1 | 2.9 | 9.9 |
| % in habitats | 344 | 9.3 | 17.4 | 64.3 | 9.0 | 100.0 |

**Additional file: Table S7.**
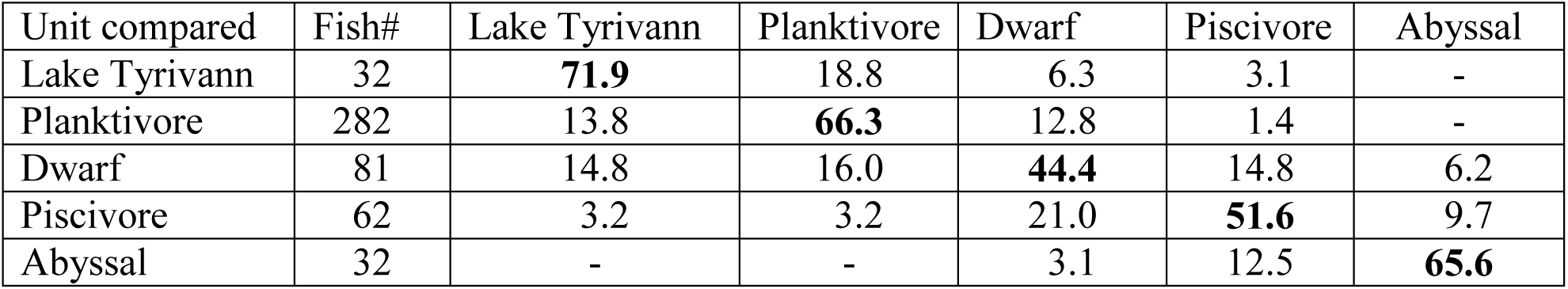
Assignment percentage based on discriminant analysis of PC-axis 1-5 for body shape in four FA-morphs in Lake Tinnsjøen compared a putative ancestor in Lake Tyrivann. The bold values denote “correct” back assignment to the original population or morph categories.

| Unit compared | Fish# | Lake Tyrivann | Planktivore | Dwarf | Piscivore | Abyssal |
| --- | --- | --- | --- | --- | --- | --- |
| Lake Tyrivann | 32 | **71.9** | 18.8 | 6.3 | 3.1 | - |
| Planktivore | 282 | 13.8 | **66.3** | 12.8 | 1.4 | - |
| Dwarf | 81 | 14.8 | 16.0 | **44.4** | 14.8 | 6.2 |
| Piscivore | 62 | 3.2 | 3.2 | 21.0 | **51.6** | 9.7 |
| Abyssal | 32 | - | - | 3.1 | 12.5 | **65.6** |

**Additional file: Table S8.** Genetic diversity measures for microsatellites summarized for the four Lake Tinnsjøen morphs and four outgroup lakes.

| Morph/<br>population | Number of<br>alleles | Private allele<br>richness | Allelic<br>richness | F<sub>IS</sub> | Heterozygosity | Gene<br>diversity |
| --- | --- | --- | --- | --- | --- | --- |
| Lake Tinnsjøen | 163 | 0.29 | 8.09 | 0.020 | 0.560 | 0.750 |
| *Planktivore* | 185 | 0.38 | 8.63 | -0.006 | 0.802 | 0.690 |
| *Dwarf* | 73 | 0.22 | 8.23 | 0.020 | 0.674 | 0.677 |
| *Piscivore* | 57 | 0.13 | 7.46 | 0.035 | 0.128 | 0.602 |
| *Abyssal* | 30 | 0.41 | 8.03 | 0.048 | 0.653 | 0.683 |
| Lake Tyrivann | 32 | 0.52 | 7.56 | -0.012 | 0.820 | 0.683 |
| Lake Vatnevatn | 32 | 0.68 | 6.02 | 0.024 | 0.160 | 0.567 |
| Lake Leirfoss | 29 | 0.69 | 7.53 | 0.118 | 0.178 | 0.761 |
| Lake Femund | 14 | 0.52 | 7.56 | -0.012 | 0.820 | 0.683 |

**Additional file: Table S9.** Output of hierarchical STRUCTURE runs using STRUCTURE-HARVESTER evaluations regarding the most likely K clusters in Lake Tinnsjøen and the four Norwegian outgroup lakes. The most likely number of K clusters were interpreted to be K=8 based on these hierarchical analyses.

|  | K | LnP(K) | Stdev LnP(K) | Ln'(K) | \|Ln''(K)\| | DeltaK |
| --- | --- | --- | --- | --- | --- | --- |
| Run 1 | 5 | -16833,62 | 2,2533 | 425,38 | 719,34 | 319,239317 |
| Run 2 | 6 | -15537,47 | 106,8824 | 481,33 | 926,71 | 8,670372 |
| Run 3 | 4 | -14525,71 | 3,8748 | 471,5 | 360,12 | 92,938191 |
| Run 4 | 5 | -16833,62 | 2,2533 | 425,38 | 719,34 | 319,239317 |
| Run 5 | 4 | -12521,01 | 85,8263 | 118,74 | NA | NA |

**Additional file: Table S10.**
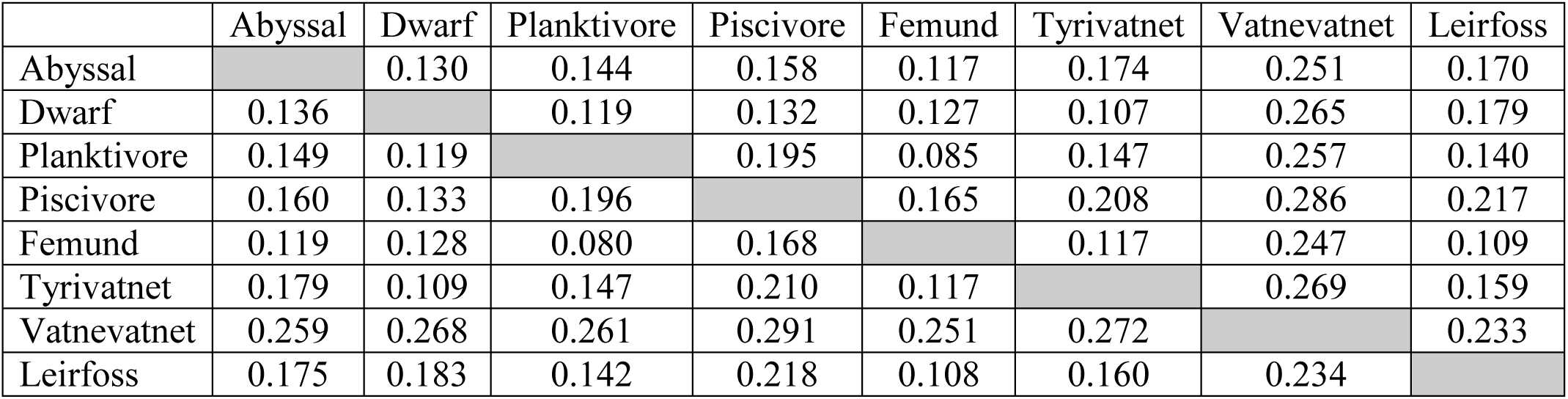
F_ST_ values for FA-morphs in Lake Tinnsjøen and four outgroup lakes. Lower diagonal; using conventional methods and upper diagonal: using ENA corrections. All comparisons are significant.

|  | Abyssal | Dwarf | Planktivore | Piscivore | Femund | Tyrivatnet | Vatnevatnet | Leirfoss |
| --- | --- | --- | --- | --- | --- | --- | --- | --- |
| Abyssal |  | 0.130 | 0.144 | 0.158 | 0.117 | 0.174 | 0.251 | 0.170 |
| Dwarf | 0.136 |  | 0.119 | 0.132 | 0.127 | 0.107 | 0.265 | 0.179 |
| Planktivore | 0.149 | 0.119 |  | 0.195 | 0.085 | 0.147 | 0.257 | 0.140 |
| Piscivore | 0.160 | 0.133 | 0.196 |  | 0.165 | 0.208 | 0.286 | 0.217 |
| Femund | 0.119 | 0.128 | 0.080 | 0.168 |  | 0.117 | 0.247 | 0.109 |
| Tyrivatnet | 0.179 | 0.109 | 0.147 | 0.210 | 0.117 |  | 0.269 | 0.159 |
| Vatnevatnet | 0.259 | 0.268 | 0.261 | 0.291 | 0.251 | 0.272 |  | 0.233 |
| Leirfoss | 0.175 | 0.183 | 0.142 | 0.218 | 0.108 | 0.160 | 0.234 |  |

**Additional file: Table S11.** F_ST_ values for the five lakes studied combining four FA-morphs in Lake Tinnsjøen into one group. Lower diagonal; using conventional methods and upper diagonal: using ENA corrections. All the comparisons are significant.

|  | Tinnsjøen | Femund | Tyrivatnet | Vatnevatnet | Leirfoss |
| --- | --- | --- | --- | --- | --- |
| Tinnsjøen |  | 0.057 | 0.101 | 0.199 | 0.113 |
| Femund | 0.057 |  | 0.117 | 0.247 | 0.109 |
| Tyrivatnet | 0.100 | 0.117 |  | 0.269 | 0.159 |
| Vatnevatnet | 0.203 | 0.251 | 0.272 |  | 0.233 |
| Leirfoss | 0.117 | 0.108 | 0.160 | 0.234 |  |

**Additional file: Table S12.** F_ST_ values for “genetically pure” (i.e. q>0.7) GA-morphs in Lake Tinnsjøen. Lower diagonal; using conventional methods and upper diagonal: using ENA corrections. All the comparisons are significant.

|  | Planktivore | Dwarf | Piscivore | Abyssal |
| --- | --- | --- | --- | --- |
| Planktivore |  | 0.087 | 0.212 | 0.130 |
| Dwarf | 0.088 |  | 0.185 | 0.126 |
| Piscivore | 0.212 | 0.185 |  | 0.201 |
| Abyssal | 0.135 | 0.131 | 0.201 |  |

**Additional file: Figure S1.**
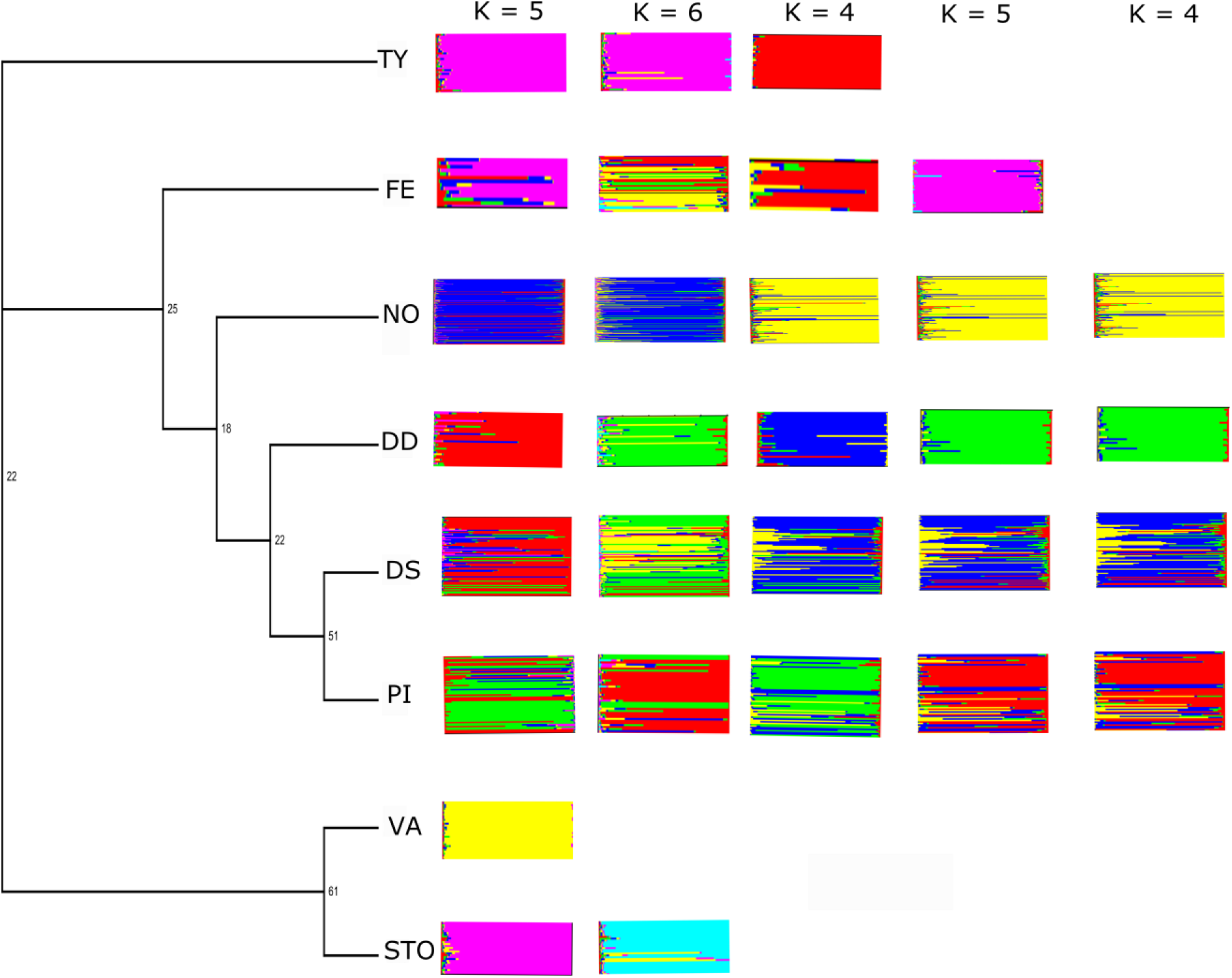
Hierarchical STRUCTURE plot using the four morphs in Lake Tinnsjøen and the four Norwegian outgroup lakes. NO=plantivore morph, DD=abyssal morph, DS=dwarf, PI=piscivore morph, while TY, FE, VA and STO are outgroup lakes in Norway.

